# Eukaryotic translation initiation factor eIF4E-5 is required for spermiogenesis in *Drosophila melanogaster*

**DOI:** 10.1101/2021.12.19.473358

**Authors:** Lisa Shao, Jaclyn M. Fingerhut, Brook L. Falk, Hong Han, Giovanna Maldonado, Yuemeng Qiao, Vincent Lee, Elizabeth Hall, Liang Chen, Gordon Polevoy, Greco Hernández, Paul Lasko, Julie A. Brill

**Author notes:** To whom correspondence should be addressed: Julie A. Brill, The Hospital for Sick Children PGCRL Building, 15^th^ Floor 686 Bay Street, Room 15.9716 Toronto, Ontario M5G 0A4, CANADA.

## Abstract

*Drosophila* sperm development is characterized by extensive post-transcriptional regulation whereby thousands of transcripts are preserved for translation during later stages. A key step in translation initiation is the binding of eukaryotic initiation factor 4E (eIF4E) to the 5’ mRNA cap. *Drosophila* has multiple paralogs of eIF4E, including four (eIF4E-3, -4, -5, and -7) that are highly expressed in the testis. Other than eIF4E-3, none of these has been characterized genetically. Here, using CRISPR/Cas9 mutagenesis, we determined that eIF4E-5 is essential for male fertility. *eIF4E-5* mutants exhibit defects during post-meiotic stages, including a fully penetrant defect in individualization, resulting in failure to produce mature sperm. eIF4E-5 protein localizes to the distal ends of elongated spermatid cysts, where it regulates non-apoptotic caspase activity during individualization by promoting local accumulation of the E3 ubiquitin ligase inhibitor Soti. *eIF4E-5* mutants also have mild defects in spermatid cyst polarization, similar to mutants affecting the cytoplasmic polyadenylation-element binding protein Orb2 and atypical protein kinase C (aPKC). Our results further extend the diversity of non-canonical eIF4Es that carry out distinct spatiotemporal roles during spermatogenesis.

**Summary Statement:** The testis-enriched translation initiation factor eIF4E-5 is needed for spermatid cyst polarization, individualization of mature sperm and male fertility in *Drosophila*.

## Introduction

Translation of mRNA into protein is frequently targeted by genetic regulatory mechanisms (Kugler and Lakso, 2009; Lin and Holt, 2007; Mingle et al., 2005; Moor et al., 2017). Protein synthesis can be divided into three steps (initiation, elongation, termination), yet much of the regulation occurs at the first step (Hershey et al., 2018; Pelletier and Sonenberg, 2019; Sonenberg and Hinnebusch, 2009). During initiation, the eukaryotic initiation factor 4F (eIF4F) cap-binding complex is recruited to the 7-methylguanylate cap located at the 5’ end of mRNAs. eIF4F is composed of a cap-binding protein (eIF4E) and an RNA helicase (eIF4A) held together by a scaffolding protein (eIF4G). eIF4G binds poly(A)-binding protein (PABP) and eIF3 to recruit the 40S ribosomal subunit. Translational repression can occur when eIF4E-binding proteins (4E-BPs) bind eIF4E to block its association with eIF4G or when eIF4E-homologous proteins (4E-HPs) bind the 5’ cap to block the recruitment of eIF4E to the mRNA. Therefore, eIF4E plays an essential role in eukaryotic cap-mediated translation initiation.

Most eukaryotic genomes encode several paralogs of eIF4E, but their roles and regulation are not well understood. Mammals have three paralogs (eIF4E1 to eIF4E3; Joshi et al., 2004): mouse eIF4E1 and eIF4E2/4EHP mutants exhibit behavioral defects (Aguilar-Valles et al., 2018; Wiebe et al., 2020). *C. elegans* has five paralogs (IFE-1 to IFE-5): IFE-3, which is most similar to mammalian eIF4E1 (*i.e.,* canonical), is essential for viability; IFE-1 is required for spermatogenesis and oocyte maturation; IFE-2 is needed for meiotic recombination; and IFE-4 is required to translate mRNAs involved in egg laying (Amiri et al., 2001; Dinkova et al., 2005; Henderson et al., 2009; Kawasaki et al., 2011; Keiper et al., 2000; Song et al., 2010). *Drosophila* has eight paralogs (eIF4E-1 to eIF4E-8/4E-HP) that bind to mRNA 5’ caps with varying affinities *in vitro* (Zuberek et al., 2016). Canonical eIF4E-1 and 4E-HP are essential for viability, and eIF4E-3 is needed for meiotic chromosome segregation and cytokinesis during spermatogenesis (Brown et al., 2014; Chintapalli et al., 2007; Ghosh and Lasko, 2015; Hernández et al., 2012). However, the roles of the remaining paralogs are uncharacterized. Whereas *eIF4E-1* and *4E-HP* mRNAs are ubiquitously expressed, *eIF4E-3, eIF4E-4, eIF4E-5* and *eIF4E-7* mRNAs are highly and specifically enriched in the testis (Graveley et al., 2011). This suggests that these paralogs may have distinct cellular or developmental roles during sperm development.

The stages of *Drosophila* male germ cell development are organized in a spatiotemporal manner (Fig. S1). The stem cell niche is located at the apical tip of the testis and germ cell development progresses toward the basal end of the testis where mature sperm will eventually exit (reviewed in Fabian and Brill 2012; Fuller 1993; Lindsley and Tokuyasu, 1980; Renkawitz-Pohl et al., 2005). Male germline stem cells divide asymmetrically to form new stem cells and differentiating daughter cells called gonialblasts. Each gonialblast undergoes four rounds of mitosis with incomplete cytokinesis to generate a cyst of 16 primary spermatocytes. After an extended period of growth and gene expression, the 16 spermatocytes undergo meiosis with incomplete cytokinesis to form a cyst of 64 interconnected haploid spermatids. A series of dramatic morphological changes convert these round spermatids into 1.85mm long mature sperm through a process called spermiogenesis. These changes include polarization of spermatid cysts relative to the long axis of the testis; assembly of flagellar axonemes that make up the sperm tails; elongation of spermatid cysts by membrane addition at the distal, growing ends; and individualization, which separates fully elongated, interconnected spermatids into individual sperm. During individualization, unneeded organelles and other cellular materials are stripped from elongated spermatid cysts by an actin-based individualization complex that moves down the length of the spermatids, forming a cystic bulge whose contents eventually pinch off into a structure called the waste bag.

Post-transcriptional regulation, possibly at the level of translation, is a crucial aspect of *Drosophila* spermatogenesis. Many genes needed post-meiotically are transcribed in primary spermatocytes and translationally repressed until later stages (Jayaramaiah-Raja and Renkawitz-Pohl, 2005; Renkawitz-Pohl et al., 2005; Santel et al., 1997; Schäfer et al., 1993; White-Cooper, 2010; White-Cooper and Caporilli, 2013; Zhao et al., 2010). In addition, a small subset of “cup” and “comet” genes, named based on their mRNA distribution, are transcribed post-meiotically, and the corresponding mRNAs localize at the growing ends of elongating spermatid cysts (Barreau et al., 2008a,b). One of these genes, *soti,* encodes a protein translated late in sperm development that regulates non-apoptotic caspase activity during individualization (Kaplan et al., 2010). Although little is known about how translation of cup and comet mRNAs is regulated, the cytoplasmic polyadenylation element binding protein (CPEB) Orb2 binds the 3’ UTR of *soti* and *f-cup,* suggesting it might regulate their translation (Xu et al., 2012). Orb2, together with atypical protein kinase C (aPKC), ensures polarization of spermatid cysts relative to the long axis of the testis (Xu et al., 2014).

Here, we show that the testis-enriched translation initiation factor eIF4E-5 is essential for male fertility, localizes to the distal ends of elongated spermatid cysts, is required for normal accumulation of Soti at the distal ends of elongated spermatids, and interacts genetically with *orb2* and *apkc* during spermatid cyst polarization. Thus, eIF4E-5 is novel player in post-transcriptional regulation during spermiogenesis.

## Results

### *eIF4E-5* is required for male fertility

*eIF4E-5* encodes a single predicted polypeptide of 232 amino acids that contains conserved residues needed to bind the mRNA cap (Asp108, Trp120, Glu121; Hernández et al. 2005). To examine the role of eIF4E-5 in *Drosophila* spermatogenesis, CRISPR/Cas9 mediated mutagenesis was used to produce *eIF4E-5* alleles with deletions in the coding sequence of the gene (Fig. 1A). The deletions in *eIF4E-5^B8a^* (12 bp), *eIF4E-5^B8b^* (3 bp) and *eIF4E-5^D19a^* (1 bp) are predicted to produce a 4-amino acid in-frame deletion, a 1-amino acid in-frame deletion, and a frame-shift resulting in a 77-amino acid truncated protein lacking the eIF4E domain, which contains the cap-binding site (Fig. 1B). All three mutations affect the conserved motif required for binding to the eIF4E-binding motif found in eIF4Gs and other eIF4E-binding proteins (His55, Pro56, Leu57; Grüner et al., 2016): *eIF4E-5^B8a^* removes Pro56 and Leu57; *eIF4E-5^B8b^* removes His55; and *eIF4E-5^D19a^* has a frameshift after His55 that removes all subsequent amino acids, replacing them with 22 amino acids from an alternate reading frame. To investigate whether levels of eIF4E-5 were reduced in these mutants, polyclonal antibodies were raised against the full-length sequence of eIF4E-5. The antibodies strongly recognized a protein at the predicted molecular weight of approximately 26.9 kDa on immunoblots of testis extracts from wild type. The level of this protein was substantially reduced in testis extracts from *eIF4E-5^B8a^* or *eIF4E-5^B8b^,* and not detectable in *eIF4E-5^D19a^* homozygous mutants (Figs 1C and S2). Thus, all three *eIF4E-5* mutations are predicted to interfere with eIF4G binding and eIF4F complex formation and result in reduced levels of eIF4E-5 protein.

**Fig. 1.**
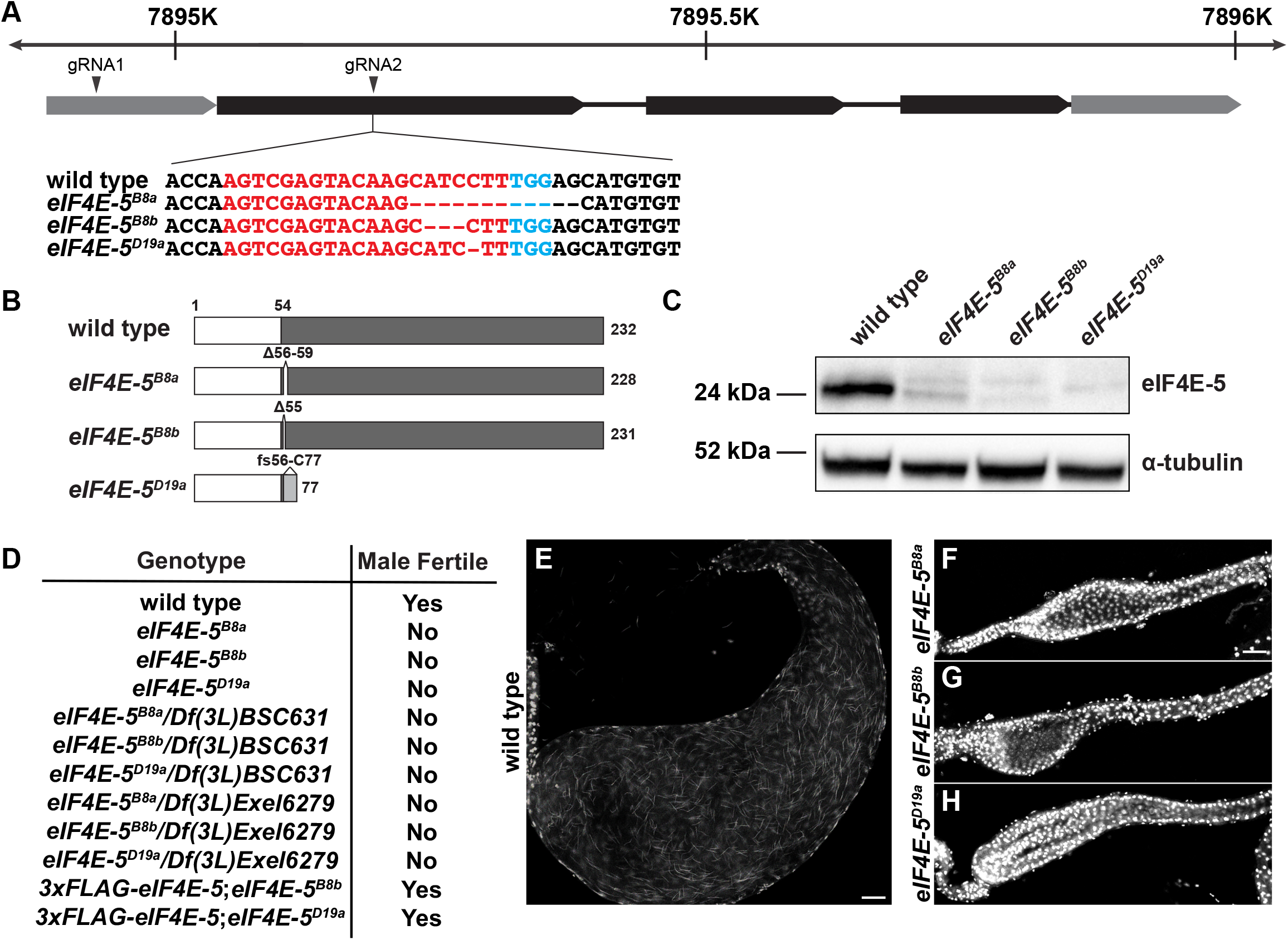
*eIF4E-5* mutants are male sterile. **(A)** CRISPR/Cas9 mutagenesis was used to generate mutants of *eIF4E-5.* Diagram showing nucleotide position of *eIF4E-5* on chromosome 3L (top). Gene structure: 5’ and 3’ UTRs (grey boxes), coding exons (black boxes), introns (lines). Locations of gRNAs in 5’UTR (gRNA1) and first exon (gRNA2) are indicated. *eIF4E-5^B8a^, eIF4E-5^B8b^* and *eIF4E-5^D19a^* mutants contain deletions in the first exon, overlapping gRNA2. Genomic sequence: protospacer adjacent motif (blue), target region of the gRNA (red), deletions (dashed lines). **(B)** Diagram showing structure of predicted wild-type (top) and mutant eIF4E-5 proteins. Wild-type eIF4E-5 has a non-conserved N-terminus (amino acids 1-54; white box) and conserved C-terminus (amino acids 54-232; dark grey box). In-frame deletions encoded by *eIF4E-5^B8a^* and *eIF4E-5^B8b^* and frameshift (fs) mutation encoded by *eIF4E-5^D19a^* (out-of-frame amino acids 56-77; light grey box) are indicated. **(C)** Immunoblot of total testis extracts probed with the indicated antibodies reveal reduced (*eIF4E-5^B8a^* and *eIF4E-5^B8b^*) or undetectable (*eIF4E-5^D19^*) levels of eIF4E-5 protein. Note that these levels correlate with the severity of the phenotypes shown in Figs 2, 3 and 7. **(D)** Fertility tests reveal that all tested combinations of *eIF4E-5* alleles and deficiencies are male sterile and that a genomic transgene expressing 3xFLAG-eIF4E-5 restores fertility to *eIF4E-5^B8b^* and *eIF4E-5^D19a^* mutant males (*eIF4E-5^B8a^* not tested). **(E-H)** Laser-scanning confocal fluorescence micrographs demonstrate the presence of needle-shaped sperm nuclei in wild-type (E) but not *eIF4E-5^B8a^* (F), *eIF4E-5^B8b^* (G) or *eIF4E-5^D19a^* (H) seminal vesicles stained with DAPI. Scale bars: 20 μm.

Males homozygous mutant for *eIF4E-5* were viable and sterile, as were male *eIF4E-5^B8a^, eIF4E-5^B8b^* and *eIF4E-5^D19a^* mutants *in trans* with two different deficiencies (*Df(3L)BSC631* and *Df(3L)Exel6279)* that uncover the *eIF4E-5* locus (Fig. 1D). A 1610 bp genomic rescue construct containing an N-terminal 3xFLAG in frame with the eIF4E-5 coding region restored male fertility in *eIF4E-5^B8b^* and *eIF4E-5^D19a^* homozygous mutants. These results confirm that the male sterility of *eIF4E-5* mutants is due to loss of eIF4E-5 function rather than a second-site mutation generated by CRISPR/Cas9 mutagenesis. Closer inspection revealed that sperm failed to accumulate in the seminal vesicles of *eIF4E-5* mutants (Fig. 1E-H) and that expression of the transgene resulted in the presence of mature, motile sperm (Fig. S3A-G). Hence, eIF4E-5 is required for male fertility.

### eIF4E-5 localizes to the distal ends of elongated spermatid cysts

To begin to decipher the requirement for *eIF4E-5* in male fertility, we examined eIF4E-5 protein distribution during sperm development. Immunostaining revealed that eIF4E-5 (yellow arrowheads) localized near the membrane skeletal protein Adducin (red arrowheads) at the distal ends of elongated spermatid cysts (Fig. 2A-A”). In contrast, elongated spermatid cysts from *eIF4E-5^B8a^, eIF4E-5^B8b^* and *eIF4E-5^D19a^* homozygotes retained Adducin localization (red arrowheads) but lacked detectible eIF4E-5 at the distal ends (Fig. 2B-D”). To confirm the localization of eIF4E-5, we examined the distribution of 3xFLAG-eIF4E-5 expressed from the rescuing transgene. Immunostaining of testes from *3xFLAG-eIF4E-5;eIF4E-5^D19a^* males with anti-FLAG and anti-eIF4E-5 antibodies revealed that these antibodies stained the same region at the distal ends of elongated spermatid cysts (Fig. 2E-E”). Moreover, testes from *eIF4E-5^D19a^* homozygotes lacked similar staining (Fig. 2F-F”), confirming that the signals were specific to eIF4E-5. In addition, immunostaining of testes from *3xFLAG-eIF4E-5;eIF4E-5^D19a^* males revealed that 3xFLAG-eIF4E-5 was present in the cytoplasm of spermatocytes and elongating spermatids (Fig. 2G-G”). In contrast, testes from *eIF4E-5^D19a^* mutants revealed non-specific staining with anti-FLAG antibodies at the tip of the testis and along a subset of elongated spermatid cysts, as well as non-specific staining with anti-eIF4E-5 antibodies in the nuclei of spermatogonia, spermatocytes and spermatids (Fig. 2F-F”,H-H”). Together, these results reveal that eIF4E-5 is expressed in early spermatocytes and persists through spermiogenesis, with enrichment at the distal ends of elongated spermatid cysts.

**Fig. 2.**
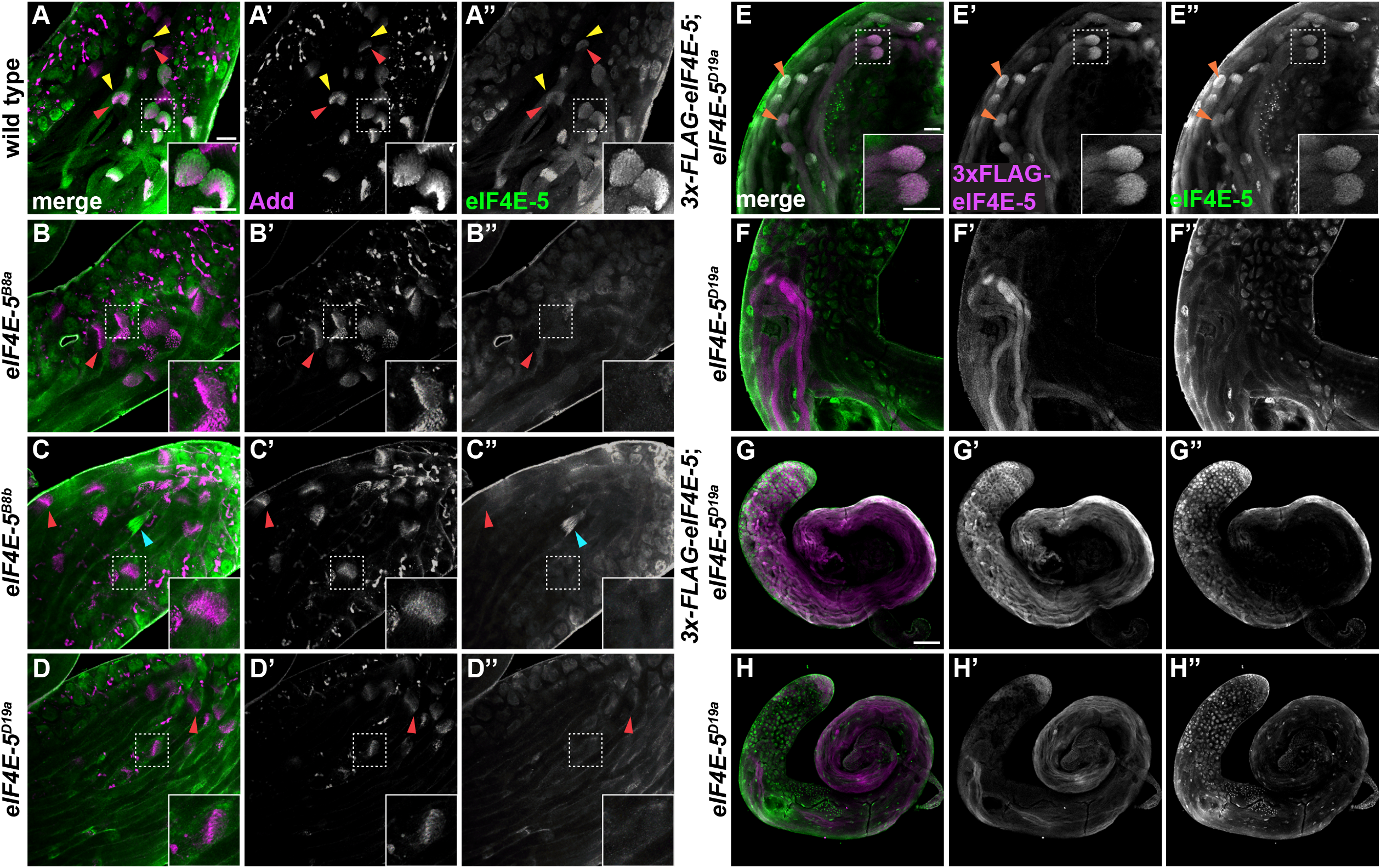
eIF4E-5 localizes to the distal ends of elongated spermatid cysts. Laser-scanning confocal fluorescence micrographs **(A-D)** Wild-type (A), *eIF4E-5^B8a^* (B), *eIF4E-5^B8b^* (C) and *eIF4E-5^D19a^* (D) whole adult testes stained for Adducin (magenta, red arrowheads) and eIF4E-5 (green, yellow arrowheads) reveals that eIF4E-5 localizes just distal to Adducin at the ends of elongated spermatid cysts in wild-type but not *eIF4E-5* mutant spermatid cysts. Note that anti-eIF4E-5 antibodies non-specifically stain individualization complexes, as shown for the mispolarized cyst in *eIF4E-5^B8b^* (C-C”, cyan arrowheads). Scale bars: 20 μm. **(E-H)** *3x-FLAG-eIF4E-5;eIF4E-5^D19a^* (E, G) and *eIF4E-5^D19a^* (F, H) whole adult testes stained for 3xFLAG-eIF4E-5 (magenta) and endogenous eIF4E-5 (green). 3xFLAG-eIF4E-5 and eIF4E-5 colocalize at the distal end of elongated spermatid cysts (E; orange arrowheads). Note non-specific staining of spermatogonia and spermatid tails with anti-FLAG antibodies and non-specific staining of nuclei in spermatogonia, spermatocytes and round spermatids with anti-eIF4E-5 antibodies (E-H”) in *eIF4E-5^D19a^* mutants. Scale bars: 20 μm (E-F), 100 μm (G-H).

### eIF4E-5 is required for spermatid individualization

To determine the cause of male sterility in *eIF4E-5* mutants, we examined testes by phase-contrast microscopy. Although overall testis morphology appeared normal, waste bags (Fig. 3A’, red arrowheads) were absent, suggesting that loss of eIF4E-5 caused defects in spermatid individualization (Fig. 3A-D’). Waste bag formation was rescued by the 3xFLAG-eIF4E-5 transgene (Fig. S3G,G’). Whole testes stained for activated (cleaved) caspase-3 revealed normal cystic bulges in wild-type testes (Fig. 3E,E’, yellow arrowheads), whereas these appeared flattened in *eIF4E-5* mutants (Fig. 3F-H’, yellow arrowheads). The mutants also displayed an elevated level of active effector caspases starting at the cystic bulges and extending further towards the distal ends of the tails (Fig. 3F’-H’), as compared to wild type (Fig. 3E’). Furthermore, unlike the synchronous movement of the 64 actin-based investment cones within caspase-positive cystic bulges in wild type (Fig. 3E”-E”’), the actin cones in *eIF4E-5* mutants appeared scattered and disorganized within the cystic bulges (Fig. 3F”-H”’). As individualization is an essential step in the formation of mature sperm, this fully penetrant phenotype is likely the cause of male infertility in *eIF4E-5* mutants.

**Fig. 3.**
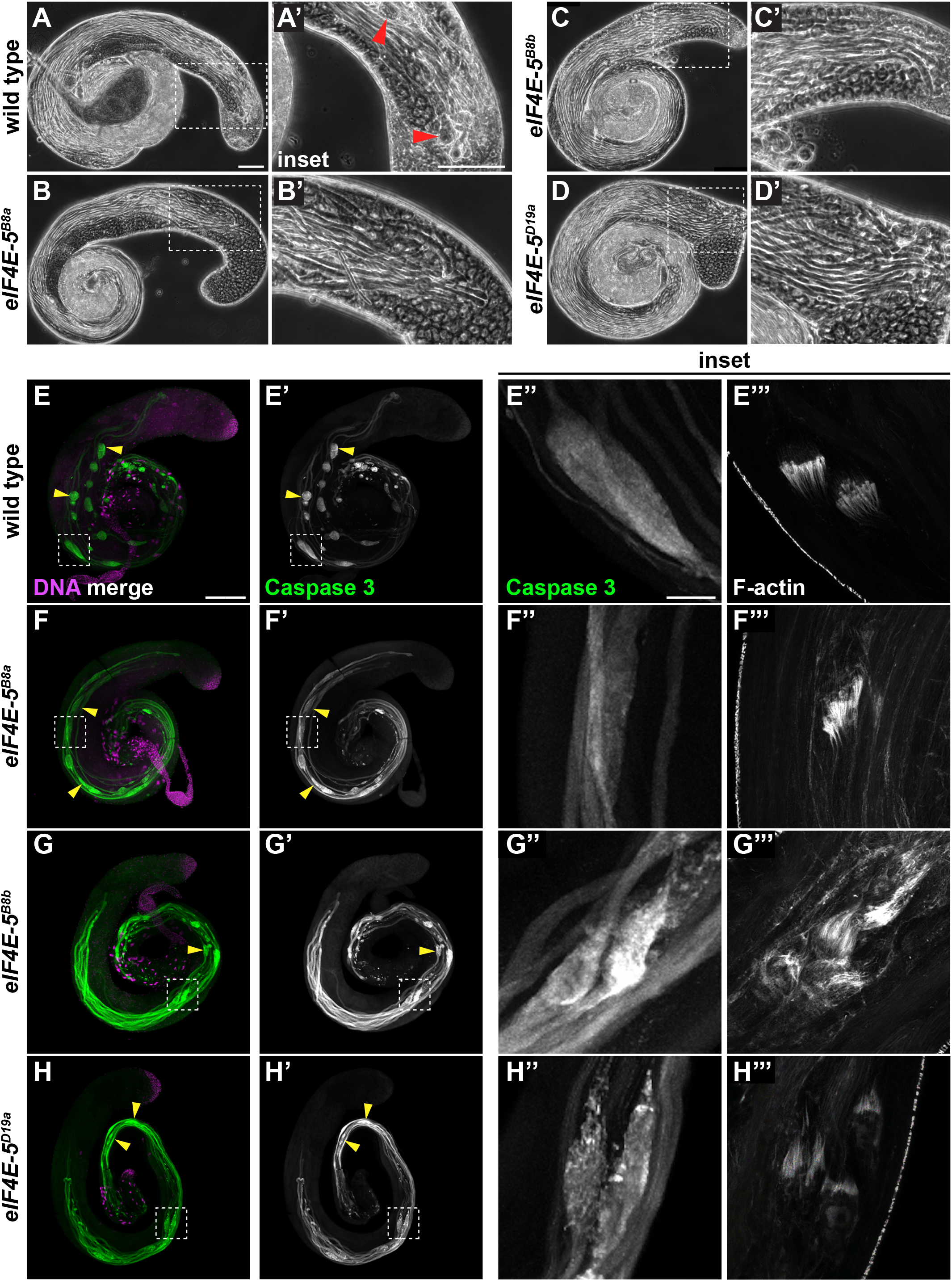
*eIF4E-5* mutants exhibit defects in individualization. **(A-D)** Phase-contrast images of 4-day old wild-type (A), *eIF4E-5^B8a^* (B), *eIF4E-5^B8b^* (C) or *eIF4E-5^D19a^* (D) testes reveal an absence of waste bags (red arrowheads) near the testis tip in *eIF4E-5* mutants. Scale bars: 100 μm. **(E-H)** Laser-scanning confocal fluorescence micrographs of wild type (E), *eIF4E-5^B8a^* (F), *eIF4E-5^B8b^* (G) or *eIF4E-5^D19a^* (H) whole adult testes stained for DNA (magenta), activated caspase-3 (green) and F-actin (shown only in insets). Activated caspase (yellow arrowheads) is restricted to cystic bulges in wild type (E) but not *eIF4E-5* mutants (F-H). Boxed areas are magnified 10-fold in insets. Groups of actin cones in individualization complexes move synchronously down the length of cysts in wild type (E”’) but become scattered prior to reaching the distal end of elongated spermatid cysts in *eIF4E-5* mutants (F”’-H”’). Brightness and contrast for Fig. 3H”’ were adjusted separately from the rest of the images for visualizing individualization complexes. Scale bars: 200 μm (whole tissue), 20 μm (insets).

### eIF4E-5 binds translational regulators 4E-BP, eIF4G-2, 4E-T and Cup

To begin to examine the biochemical properties of eIF4E-5, we tested whether eIF4E-5 binds known eIF4E binding proteins, including eIF4G and eIF4G-2, which have roles in sperm development and male fertility (Baker and Fuller 2007; Franklin-Dumont et al., 2007; Ghosh and Lasko 2015). We performed yeast two-hybrid assays using eIF4E-5 as “bait” and constructs containing 4E-BP (Miron et al. 2001), eIF4G (Hernández et al. 1998), eIF4G-2 (Baker and Fuller 2007; Franklin-Dumont et al., 2007; Ghosh and Lasko 2015), 4E-T (Kamenska et al. 2014), Cup (Nelson et al, 2004; Zappavigna, et al., 2004) and GIGYF (Russica et al., 2019) as “prey” (Fig. S4). Positive interactions were detected between eIF4E-5 and 4E-BP, eIF4G-2, 4E-T, and Cup, but not eIF4G. The lack of interaction between eIF4E-5 and eIF4G was unexpected, as we previously detected a weak interaction of these proteins in a more sensitive yeast two-hybrid assay (Hernández et al., 2005) and in fluorescent binding assays with a short eIF4G peptide containing the eIF4E-binding motif (Zuberek et al., 2016). In addition, unlike binding of eIF4E-5 to 4E-BP, 4E-T and Cup, binding to eIF4G-2 was sensitive to more stringent selection conditions, indicating that eIF4E-5 may preferentially bind inhibitors of translation rather than translational activators. Overall, these results suggest eIF4E-5 may bind known translational regulators in various complexes during sperm development.

### eIF4E-5 is required for localized accumulation of Soti protein

As eIF4E-5 is essential for spermatid individualization and localizes at the distal ends of elongated spermatid cysts, we sought potential eIF4E-5 translational targets that show similar localization during spermiogenesis. Among the cup and comet genes, *soti* encodes a testis-specific E3 ubiquitin ligase inhibitor that controls caspase activation and is concentrated at the distal ends of elongated spermatid cysts where its mRNA is found (Barreau et al., 2008a; Kaplan et al., 2010). Because of Soti’s role in individualization and similar localization of the Soti and eIF4E-5 proteins, we investigated whether Soti protein distribution was affected in *eIF4E-5* mutants. In wild type, Soti was concentrated at the distal ends of elongated spermatid cysts (Fig. 4A, yellow arrowheads), where it colocalized with the membrane skeletal protein Adducin (Fig. 4D, red arrowheads; Hime et al., 1996). Soti protein levels were reduced in *eIF4E-5^D19a^* mutants (Fig. 4B,E), and normal levels of Soti protein were rescued by expression of the 3xFLAG-eIF4E-5 genomic transgene (Fig. 4C, yellow arrowheads). Single molecule RNA florescent *in situ* hybridization (smFISH) revealed that loss of *eIF4E-5* did not alter *soti* mRNA expression or localization (Fig. S5A-F). These results demonstrate that eIF4E-5 is required for localized accumulation of Soti protein at the distal ends of elongated spermatid cysts.

**Fig. 4.**
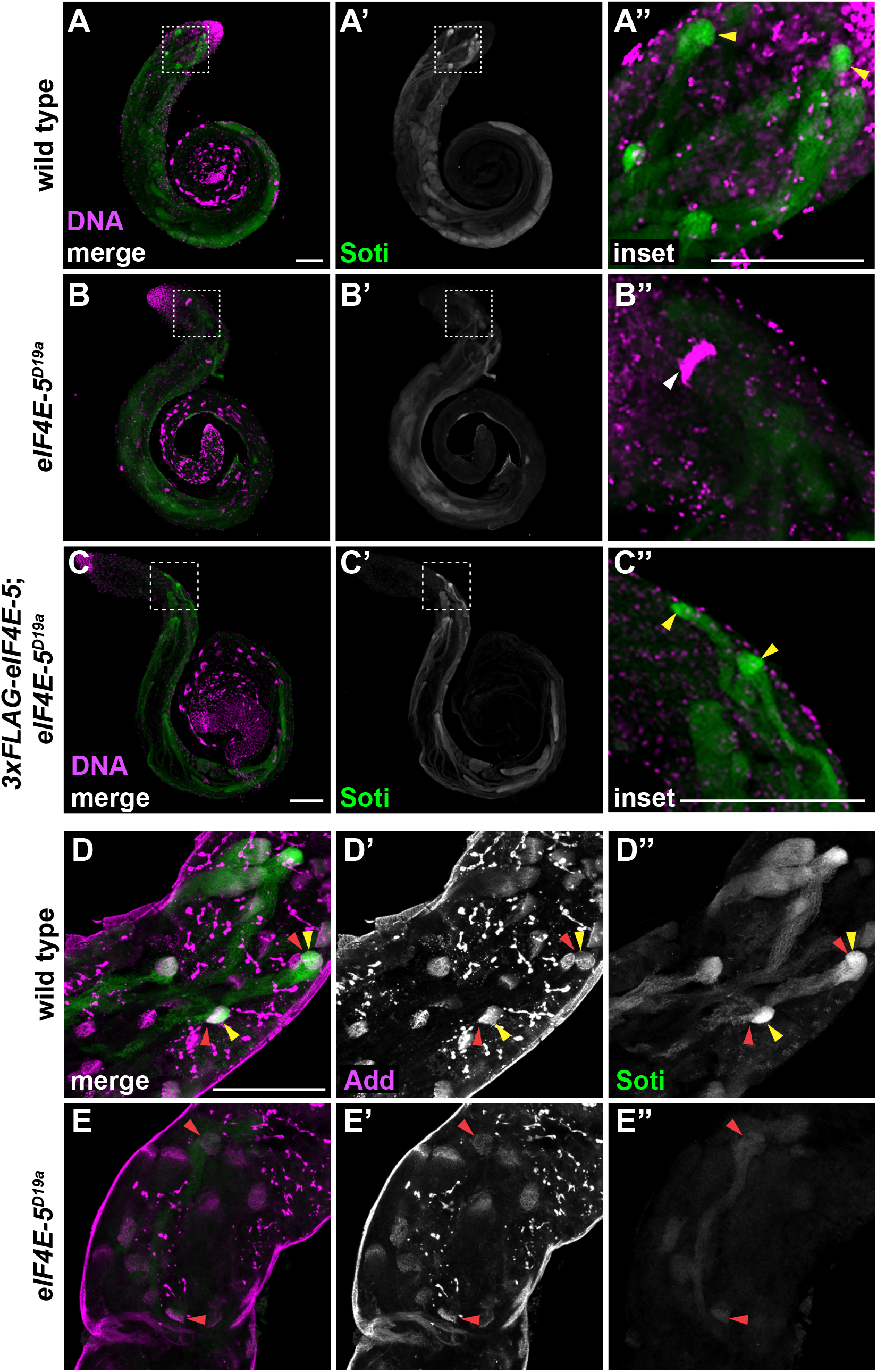
eIF4E-5 is required for localized accumulation of Soti, a caspase inhibitor. Laser-scanning confocal fluorescence micrographs. **(A-C)** Wild-type (A), *eIF4E-5^D19a^* (B) and *3xFLAG-eIF4E-5;eIF4E-5^D19a^* rescue (C) whole adult testes stained for DNA (magenta) and Soti (green). Boxed areas are magnified 4 to 5-fold in insets. Soti is enriched at distal ends of elongated spermatid cysts in wild type and rescue but not *eIF4E-5^D19a^* (yellow arrowheads). Note presence of nuclei from mispolarized spermatid cyst in *eIF4E-5^D19a^* (B”, white arrowhead). Scale bars: 100 μm. **(D-E)** Tip regions of testes stained for Soti (green, yellow arrowheads) and Adducin (magenta, red arrowheads). Soti localizes near Adducin at the distal ends of elongated spermatid cysts in wild-type (C-C”) but is reduced in *eIF4E-5^D19a^* (D-D”) testes. Scale bar: 100 μm.

Because of their similar distribution in elongated spermatid cysts, we examined whether eIF4E-5 colocalizes with *soti* mRNA or Soti protein. smFISH of *soti* mRNA combined with immunofluorescence of 3xFLAG-eIF4E-5 protein (Fig. 5A-A”) or endogenous eIF4E-5 (Fig. S5G-G”) revealed that *soti* mRNA (cysts with dashed outlines) and eIF4E-5 protein (red arrowheads) did not colocalize and were concentrated at the distal ends of different elongated spermatid cysts. In addition, smFISH showed that *eIF4E-5* mRNA was diffusely cytoplasmic starting in primary spermatocytes and did not colocalize with *soti* mRNA or become concentrated at the distal ends of spermatid cysts (Fig. S5H-J). In contrast, immunofluorescence revealed colocalization of 3xFLAG-eIF4E-5 and Soti protein (Fig. 5B-B”), with some cysts exhibiting higher (Fig. 5Bi), similar (Fig. 5Bii) or lower (Fig. 5Biii) levels of eIF4E-5 relative to Soti at the distal ends the elongated spermatid cysts. These results suggest either that eIF4E-5 promotes Soti translation concomitant with *soti* mRNA degradation or that eIF4E-5 affects accumulation of Soti at the distal ends of elongated spermatid cysts by an alternate mechanism.

**Fig. 5.**
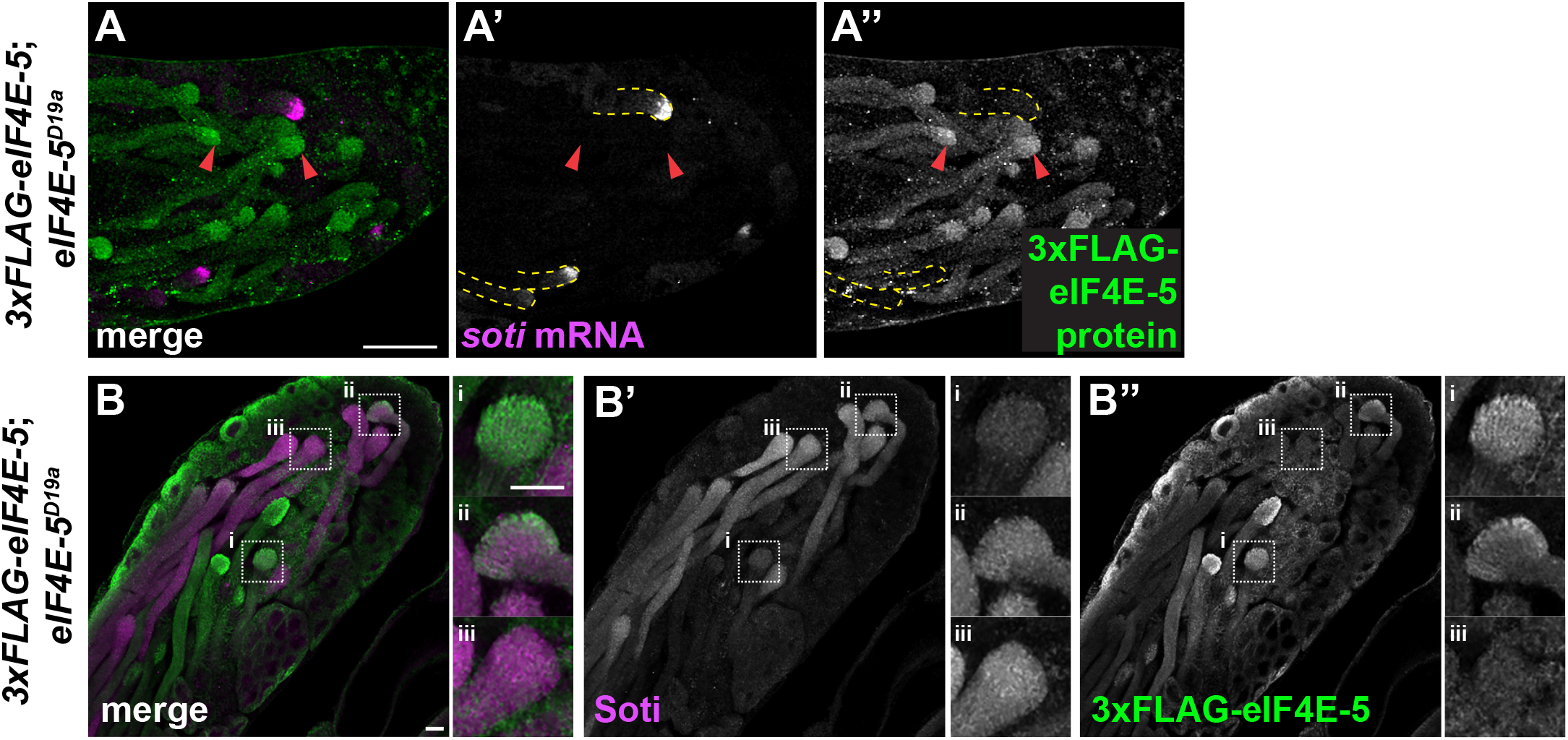
eIF4E-5 colocalizes with Soti protein but not *soti* mRNA at the distal ends of elongated spermatid cysts. Laser-scanning confocal fluorescence micrographs. **(A)** *3xFLAG-eIF4E-5;eIF4E-5^D19a^* adult testis probed for *soti* mRNA (magenta; in cysts outline by dotted yellow line) and stained for 3xFLAG-eIF4E-5 protein (green; red arrowheads). Scale bar: 50 μm. **(B)** *3xFLAG-eIF4E-5;eIF4E-5^D19a^* adult testis stained for Soti protein (magenta) and 3xFLAG-eIF4E-5 protein (green) reveals varying extents of eIF4E-5 and Soti colocalization at the distal ends of elongated spermatid cysts (i-iii). Boxed areas are magnified 3-fold in insets. Scale bar: 10 μm.

### eIF4E-5 is dispensable for translation of axonemal dyneins

Because flagellar axoneme assembly occurs at the distal ends of elongating spermatids and proper axoneme assembly is required for individualization (Fabian and Brill, 2012; Gottardo et al., 2013; Riparbelli et al., 2012; Tokuyasu, 1975), we examined whether eIF4E-5 is needed for the translation of axonemal proteins. Transcripts encoding the testis-specific axonemal dynein heavy chain Kl-3 localize at the distal ends of spermatid cysts (Fingerhut and Yamashita, 2020). We used smFISH to test whether *soti* and *kl-3* mRNAs colocalize in the same cysts and found that although both transcripts were present at the distal ends of early elongating spermatid cysts, they did not colocalize (Fig. 6A). In addition, *soti* mRNA was present at the distal ends of late elongating spermatid cysts at stages when *kl-3* mRNA was no longer enriched (Fig. 6B,C). Similar to *soti* mRNA, *kl-3* mRNA (cysts with dashed outlines) did not colocalize with eIF4E-5 protein (red arrowheads) at the distal ends of elongating cysts (Fig. 6D-F). Immunoblotting of endogenously tagged Kl-3 3xFLAG revealed that Kl-3 protein levels were unaffected in *eIF4E-5* mutants (Figs 6G and S6). Hence, *soti* and *kl-3* transcript localization appears to be distinct, and the translational regulation involved in axoneme assembly is likely independent of eIF4E-5.

**Fig. 6.**
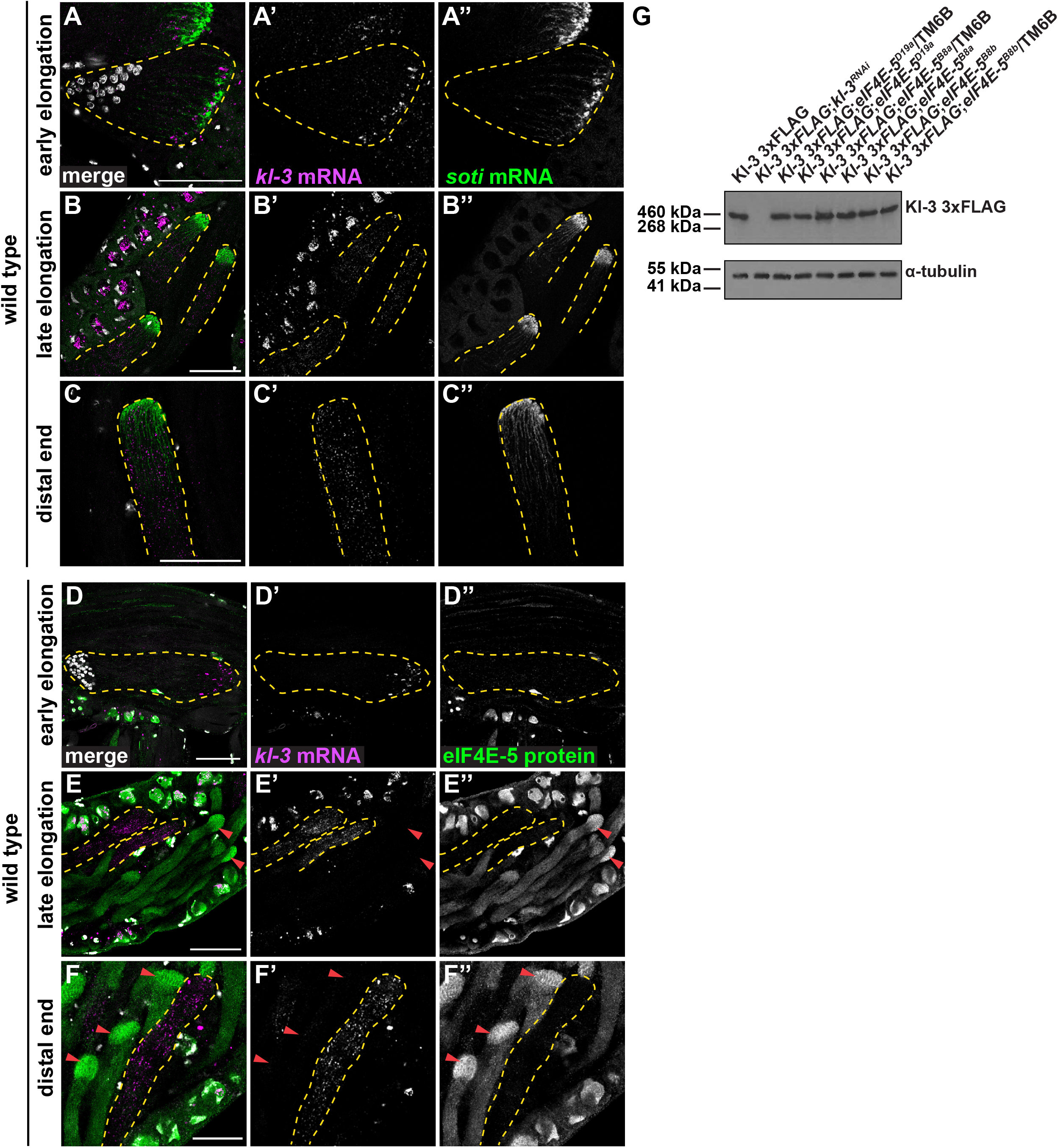
Accumulation of axonemal dynein Kl-3 is independent of *eIF4E-5*. **(A-F)** Laser-scanning confocal fluorescence micrographs. **(A-C)** Wild-type whole adult testes probed for *kl-3* mRNA (magenta) and *soti* mRNA (green) and stained for DNA (white). *kl-3* mRNA (cysts outlined by dotted yellow lines) does not colocalize with *soti* mRNA at the distal end of elongating spermatid cysts. Scale bars: 50 μm. **(D-F)** Wild-type whole adult testes probed for *kl-3* mRNA (magenta) and stained for 3xFLAG-eIF4E-5 protein (green) and DNA (white). *kl-3* mRNA (cysts outlined by dotted yellow lines) does not colocalize with eIF4E-5 protein (red arrowheads) at the distal ends of early or late elongating spermatid cysts. Scale bars: 50 μm. **(G)** Immunoblots of whole testis extracts revealing Kl-3 3xFLAG levels in the indicated genotypes expressing endogenously tagged Kl-3 3xFLAG. Kl-3 3xFLAG protein levels are unaffected in *eIF4E-5* mutants.

**Fig. 7.**
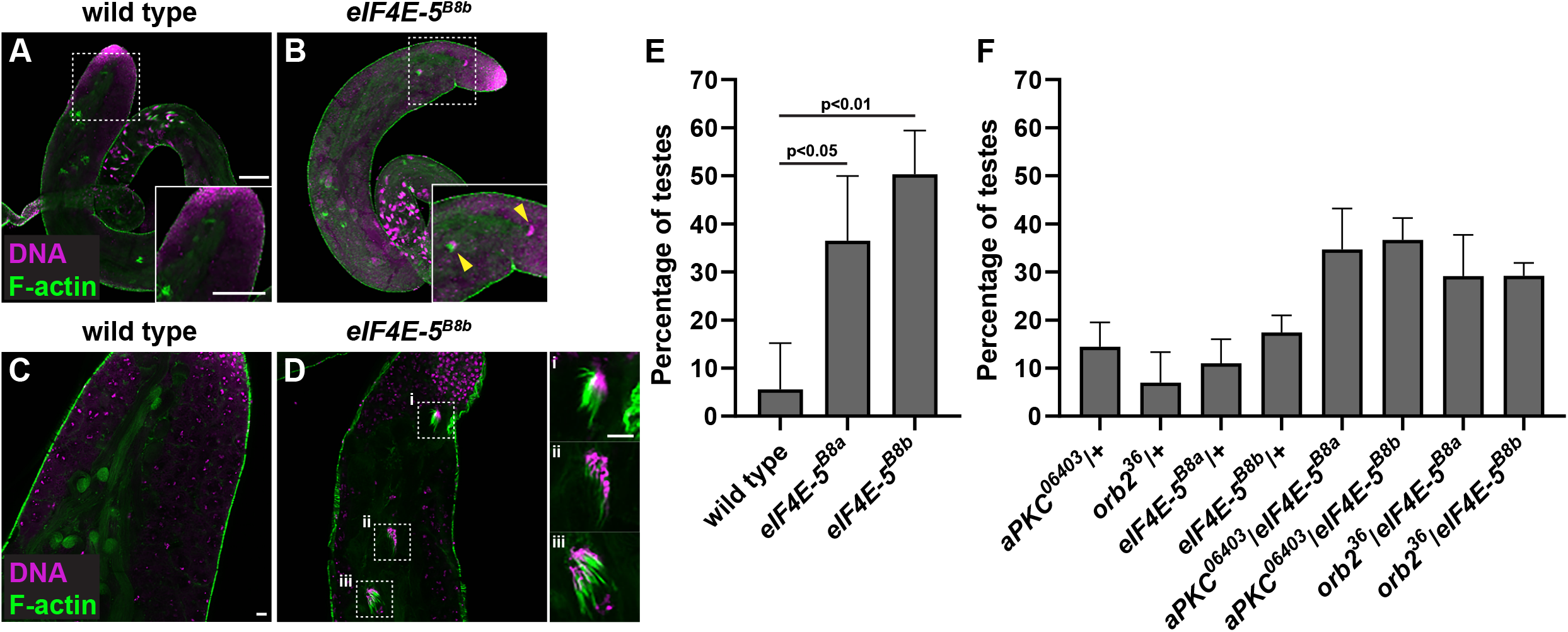
eIF4E-5 acts with Orb2 and aPKC to regulate spermatid cyst polarity. **(A-D)** Laser-scanning confocal fluorescence micrographs of wild-type (A, C) or *eIF4E-5^B8b^* (B, D) whole adult testes stained for DNA (magenta) and F-actin (green). Nuclei in elongated spermatid cysts orient towards the basal end in wild-type testes, whereas occasional clusters of nuclei orient towards the testis tip in *eIF4E-5^B8b^* mutants (B, D). Scale bars: 100 μm. **(E-F)** Percentage of testes that have at least one cluster of spermatid nuclei found at the tip instead of the basal end of the testes. Error bars show standard deviation based on three sets of experiments. Student *t*-tests were unpaired. **(E)** The percentage of testes with misoriented spermatid cysts was significantly higher in homozygous *eIF4E-5* mutants as compared to wild type: *eIF4E-5^B8a^* (*p*<0.05) and *eIF4E-5^B8b^* (*p*<0.01). Number of testes scored for each genotype, from left to right: wild type = 17; *eIF4E-5^B8a^* = 32; *eIF4E-5^B8b^* = 25. **(F)** The percentage of testes with misoriented spermatid cysts was significantly higher in each of the four transheterozygotes relative to their respective heterozygous controls: *aPKC^06403^*/+; *eIF4E-5^B8a^/+* and *aPKC^06403^*/+ (*p*<0.05) and *eIF4E-5^B8a^/+* (*p*<0.05), *aPKC^06403^/+; eIF4E-5^B8b^/+* and *aPKC^06403^/+* (*p*<0.01) and *eIF4E-5^B8b^/+* (*p*<0.01), *orb2^36^/eIF4E-5^B8a^* and *orb2^36^/+* (*p*<0.05) and *eIF4E-5^B8a^/+ (p<0.05), orb2^36^/eIF4E-5^B8b^* and *orb2^36^/+* (*p*<0.01) and *eIF4E-5^B8b^/+* (*p*=0.01). Number of testes scored for each genotype, from left to right: *aPKC^06403^*/+ = 30, *orb2^36^*/+ = 28, *eIF4E-5^B8a^*/+ = 31, *eIF4E-5^B8b^*/+ = 72, *aPKC^06403^*/+; *eIF4E-5^B8a^/+* = 145, *aPKC^06403^/+; eIF4E-5^B8b^/+* = 108, *orb2^36^/eIF4E-5^B8a^* = 117, *orb2^36^*/*eIF4E-5^B8b^* = 125.

### eIF4E-5 affects polarization of spermatid cysts

In addition to defects in individualization, *eIF4E-5* mutants exhibited spermatid cysts that were mispolarized relative to the long axis of the testis (Fig. 7A-D). In wild-type testes, spermatid cysts with elongated nuclei (~50 cysts/testis; Zhou et al., 2011) were oriented such that all 64 nuclei within a cyst point towards the basal end of the testis and the tails point towards the tip (Fig. 7A,C). In contrast, in *eIF4E-5* mutants, a small subset of cysts (at most three) was mispolarized relative to the long axis of the testis, such that clusters of elongated nuclei faced the tip (Fig. 7B, inset, yellow arrowheads and Fig. 7D, insets; see also Fig. 2C,C”, cyan arrowheads). Whereas mispolarized cysts were rarely seen in wild-type testes (5%), 30-50% of *eIF4E-5* mutant testes exhibited this phenotype (*p*<0.05), which appeared to be more common in *eIF4E-5^B8b^* mutants than *eIF4E-5^B8a^* mutants (Fig. 7E). The cyst mispolarization phenotype was also observed but not quantified in *eIF4E-5^D19a^* mutants (Fig. 4B”, white arrowhead). However, because nearly all spermatid cysts are oriented properly in testes from *eIF4E-5* mutants, this defect is likely not the cause of the observed fertility defects.

### eIF4E-5 acts with Orb2 and aPKC to promote spermatid cyst polarization

The cyst polarization defect in *eIF4E-5* mutants was reminiscent of phenotypes observed in mutants for the cell polarity regulator atypical protein kinase C (aPKC) and the cytoplasmic polyadenylation-element binding protein (CPEB) Orb2 (Xu et al., 2014). Heterozygosity for a null allele of either *aPKC* (*aPKC^k06403^*) or *orb2* (*orb2^36^*) leads to mispolarized spermatid cysts. To determine whether eIF4E-5 acts in the same manner as aPKC and Orb2, we examined testes from males transheterozygous for *eIF4E-5^B8a^* or *eIF4E-5^B8b^* and *aPKC^k06403^* or *orb2^36^* (Fig. 7F). In comparison to the single heterozygous mutants, which exhibited low levels of cyst mispolarization (7-17% of testes), this phenotype was enhanced in the transheterozygotes (29-37% testes) (*p*<0.05). These genetic results suggest that eIF4E-5 has at least a partially overlapping role with Orb2 and aPKC in controlling spermatid cyst polarization.

## Discussion

### eIF4E-5 is required for spermiogenesis and male fertility

Our data indicate that eIF4E-5 is essential for *Drosophila* male fertility and needed for faithful polarization of spermatid cysts and individualization of spermatids to form mature sperm. These post-meiotic defects in *eIF4E-5* mutants are distinct from the earlier defects observed in *eIF4E-3* mutants (Hernández et al., 2012), demonstrating that two of the four testis-enriched eIF4Es have distinct spatiotemporal roles during *Drosophila* spermatogenesis. eIF4E-3 is most highly enriched in primary spermatocytes, where it is needed for meiotic chromosome segregation and cytokinesis (Hernández et al., 2012). In contrast, eIF4E-5 concentrates at the distal end of elongated spermatid cysts, a site important for regulating individualization. Although both eIF4E-3 and eIF4E-5 are transcribed in primary spermatocytes, enrichment of these eIF4E proteins coincides with the stages they regulate, supporting the idea that the evolutionary expansion of these non-canonical testis-enriched eIF4Es allows for distinct roles and regulation during spermatogenesis. Indeed, complementation experiments revealed that eIF4E-1 and eIF4E-3, but not eIF4E-5, were able to rescue growth of *Saccharomyces cerevisiae* lacking its only eIF4E paralog, Cdc33, suggesting that eIF4E-1/-3 and eIF4E-5 likely have different activities or require distinct binding partners *in vivo.* Sorting out the corresponding mechanisms will provide a better understanding of post-transcriptional regulation in *Drosophila* sperm development.

### eIF4E-5 is needed for spermatid individualization and localized Soti protein accumulation

Non-apoptotic caspase activity is needed for progression of actin cones and degradation of unneeded organelles during individualization (Arama *et al.*, 2003; Huh *et al.*, 2004; Arama *et al.*, 2007; Muro *et al.*, 2006). In elongated spermatid cysts, caspases are initially activated at the nuclear end, where actin cones form, and repressed along the spermatid tails. As the actin cones move away from the nuclei, the peak of caspase activity remains associated with the cystic bulge. This localized caspase activity allows for controlled degradation of organelles, and disruption of this activity leads to failure of individualization, characterized by scattered actin cones. In this study, we show that localized caspase activation is disrupted and actin cones are scattered in individualizing spermatids of *eIF4E-5* mutants. These results suggest that eIF4E-5 post-transcriptionally regulates non-apoptotic caspase activity during spermiogenesis.

Caspase activity is regulated during individualization by the inhibitor of apoptosis (IAP)-like protein dBruce. dBruce is ubiquitinated by the cullin E3 ubiquitin ligase complex CRL3 and degraded at the nuclear end, allowing caspase activation (Arama *et al.*, 2007). At the distal end, Soti binds and prevents CRL3 from binding dBruce and promoting its degradation (Kaplan *et al.*, 2010). In *soti* mutants, CRL3 is activity leads to dBruce destruction and caspase activation along the entire length of the spermatid cysts, inhibiting proper individualization. Here, we show that localized accumulation of Soti protein is reduced at the distal end of elongated spermatids in *eIF4E-5* mutants, which exhibit defects in individualization that resemble *soti* mutants.

Although Soti protein and eIF4E-5 colocalize at the distal ends of elongated spermatid cysts, *soti* mRNA and eIF4E-5 do not. One possibility is that Soti translation depends on eIF4E-5 and that *soti* mRNA is degraded in a co-translational manner, as seen for many transcripts in yeast and mammals (Pelechano et al., 2015; Tuck et al., 2020). However, this seems unlikely, as some Soti protein remains present in *eIF4E-5* mutants. Another possibility is that one of the other testis eIF4Es might be able to translate *soti,* but with less efficiency than eIF4E-5, resulting in production of a smaller amount of Soti protein. Yet another explanation could be that eIF4E-5 regulates Soti translation indirectly or that it regulates other mRNA targets that are needed for individualization. In addition to *soti,* many transcripts have been identified that are post-meiotically transcribed and accumulate at the distal ends of elongating spermatid cysts in cup or comet patterns (Rathke et al., 2007; Barreau et al., 2008a,b; Vibranovski et al., 2009, 2010). It is possible that one or more of these transcripts is needed for individualization and requires eIF4E-5 for its localization or localized translation. Alternatively, eIF4E-5 might promote Soti accumulation independent of any role it may have in mRNA regulation. Thus, although our results reveal a novel requirement for eIF4E-5 in promoting regulation of non-apoptotic caspase activity during *Drosophila* spermatogenesis, its mechanism of action remains obscure. Distinguishing among these possibilities will be the subject of future studies.

### eIF4E-5 acts with aPKC and Orb2 to regulate spermatid cyst polarization

For successful transfer of mature sperm to the seminal vesicle, spermatid cysts must polarize such that nuclei face the basal end of the testis, and the tails point towards the tip. aPKC and Orb2 are involved in polarization of the cysts; heterozygous *aPKC* or *orb2* mutants exhibit bundles of 64 spermatids whose polarity is reversed relative to the long axis of the tissue (Xu *et al.*, 2014). Orb2 ensures localization and localized translation of *aPKC* mRNA to establish spermatid cyst polarity, and transheterozygotes of *aPKC* and *orb2* have a more severe defect than heterozygotes alone (Xu *et al.*, 2014). Here, we show that *eIF4E-5* mutants exhibit the same polarization defect as *aPKC* and *orb2* mutants, with a subset of spermatid cysts pointing towards the wrong end of the testis. Transheterozygous mutants of *eIF4E-5* and *aPKC* or *orb2* have a more severe polarity defect than the heterozygous mutants alone, suggesting that eIF4E-5, aPKC and Orb2 might act in the same pathway to establish spermatid cyst orientation. aPKC protein localizes at the growing ends of elongating spermatids where its mRNA is also found (Xu *et al.*, 2014), raising the possibility that *aPKC* mRNA or protein accumulation might be regulated by eIF4E-5.

In addition to localized translation of *aPKC* in spermatids, there is evidence that polarity proteins are post-transcriptionally regulated in different cell contexts (Barr *et al.*, 2016). For example, *Par-3* mRNA is locally translated for axonal outgrowth in embryonic rat neurons (Hengst *et al.*, 2009; Macara *et al.*, 2009). This raises the possibility that polarity proteins other than aPKC are similarly regulated in spermatids. Xu *et al.* (2014) showed that Bazooka, Dlg and Par-1 do not localize at the growing ends of spermatid cysts, suggesting they are unlikely targets of post-transcriptional regulation. However, the authors describe a subset of cysts as mispolarized in *par-6* heterozygotes. Thus, Par-6 might also act with eIF4E-5, aPKC and Orb2 to control spermatid cyst polarization, and that its transcript could be a potential target of eIF4E-5 regulation. Our results add to the existing literature that post-transcriptional regulation plays an important role in cyst polarization during *Drosophila* spermatogenesis and indicate that eIF4E-5 participates in this process.

### Regulation of eIF4E-5 during spermiogenesis

Although the relationship of eIF4E-5 to aPKC translation is unclear, our data show that Soti translation does not rely solely on eIF4E-5. In addition, eIF4E-5 is dispensable for translation of at least one other transcript found at the distal ends of spermatid cysts, *kl-3.* Indeed, it appears that eIF4E-5 accumulates in elongated spermatids at a later stage than *soti* and *kl-3* mRNAs. Because Kl-3 is required for construction of the flagellar axoneme during elongation, and *eIF4E-5* is required for individualization at a later stage, eIF4E-5 localization to the distal end likely initiates around the time spermatids become fully elongated. Because eIF4E-5 protein is present in the cytoplasm of male germ cells in primary spermatocytes and spermatids, and its transcript also shows a diffuse localization at these stages, it appears unlikely that eIF4E-5 itself is post-transcriptionally regulated. Thus, it remains unclear how eIF4E-5 protein becomes concentrated at the distal end. Perhaps the mRNA localization mechanism involved in transporting the cup and comet transcripts from nuclei also regulates localization of eIF4E-5 protein. Alternatively, it is possible that local translation of eIF4E-5 depends on translational machinery that is present at the distal end. It will be of interest to identify the factors needed for mRNA transport and translation at the distal ends of elongated spermatid cysts.

Our results indicate that eIF4E-5 directly binds several known eIF4E binding partners (4E-BP, eIF4G-2, 4E-T, Cup) and might act in the same pathway as Orb2. As all of these proteins regulate mRNA translation (Miron et al. 2001; Kamenska et al. 2014; Nelson et al, 2004; Zappavignia, et al., 2004; Baker and Fuller 2007; Franklin-Dumont et al., 2007), interactions with these proteins may help facilitate or repress translational activity of eIF4E-5 and its target mRNAs at different stages of spermiogenesis. Although canonical eIF4E-1 and testis-specific eIF4E-3 both associate with canonical eIF4G and testis-specific eIF4G-2 (Ghosh and Lasko, 2015), eIF4E-5 shows a stronger interaction with eIF4G-2 than eIF4G. Thus, an eIF4F complex formed by eIF4E-5 and eIF4G-2 might target transcripts for translation *in vivo*.

## Conclusion

Here, we show that the testis-specific *Drosophila* eIF4E paralog eIF4E-5 is essential for male fertility. Loss of eIF4E-5 disrupts localized accumulation of the caspase inhibitor Soti during individualization and hence regulated activation of Caspase-3. In addition, eIF4E-5 acts with Orb2 and aPKC to promote spermatid cyst polarization. Our study provides evidence of localized post-transcriptional regulation by eIF4E-5 during two developmental stages of *Drosophila* spermatogenesis. Future experiments will reveal the mechanism by which eIF4E-5 acts to promote male fertility.

Although there are apparent differences between *Drosophila* and human spermatogenesis, activation of pro-apoptotic proteins without causing the death of the entire cell is also used to eliminate cytoplasmic components during terminal differentiation of mammalian spermatids (Shaha et al., 2010). Because one known cause of human male infertility is incomplete extrusion of cytoplasm (Rengan et al., 2012), it would be of interest to know if there is a similar role for post-transcriptional regulation in promoting this aspect of male fertility in humans.

## Materials and Methods

### Fly strains and husbandry

Flies were raised on standard cornmeal molasses agar at 25°C (Ashburner, 1990). *w^1118^* was used as the experimental control. *w^1118^*; *PBac{vas-Cas9}* (Bloomington Drosophila Stock Center (BDSC) #56552, Bloomington, USA) was used to generate *eIF4E-5* CRISPR/Cas9 mutants. *y^l^ M{3xP3-RFP-3xP3-GFP-vas-int.DM}ZH-2A w*; P{CaryP}attP40* (BestGene Inc.) contains a second chromosome attP site (25C6) and was used to generate 3xFLAG-eIF4E-5 transgenic lines. Double-balancer stock *w^1118^*; Sco/CyO; MKRS/TM6B was used for balancing mutants. *eIF4E-5* alleles were examined *in trans* to chromosomal deletions Df(3L)BSC631 (BDSC #25722) and Df(3L)Exel6279 (BDSC #7745) lacking the entire *eIF4E-5* coding region. *Kl-3^3xFLAG^* carries a 3xFLAG tag at the endogenous C-terminus of the *kl-3* coding region, generated by CRISPR/Cas9 mediated knock-in (Fingerhut et al., 2019). *UAS-kl-3^TriP.HMC03546^* (BDSC #53317) expresses dsRNA for RNAi directed against *kl-3* under UAS control (Perkins et al., 2015). *aPKC^k06403^* (BDSC #10622) carries a *P{lacW}* insertion between two promoters in the third intron, resulting in a loss of function allele (Xu et al., 2014). *Orb2^36^* (BDSC #58479) carries a deletion of the Orb2 coding region generated through FRT-mediated recombination between two flanking progenitor insertions of *PBac{WH}CG43783^f04965^* and *PBac{WH}orb2^f01556^* (Xu et al., 2012).

Fertility tests were performed by mating individual males of each genotype with five virgin *w^1118^* females at 25°C. After 5 days, crosses were observed for the presence of progeny.

### Generation of *eIF4E-5* mutant strains

gRNAs targeting two different regions in *eIF4E-5* with no predicted off-targets were selected using the CRISPR Optimal Target Finder (http://targetfinder.flycrispr.neuro.brown.edu; Gratz et al., 2014): 5’-GAATTTTGTCGCGATTCGAG-3’ (gRNA1) and 5’-GAGTCGAGTACAAGCATCCTT-3’ (gRNA2). The two selected gRNAs were cloned into pCFD4d under two promoters, U6-1 and U6-3 (Addgene plasmid #83954, Watertown, USA; Ge et al., 2016). pCFD4d was digested with *BbsI* (New England Biolabs, R3539L, Waltham, USA) and gel purified. Inserts were generated by PCR using the following primers and pCFD4d as a template: 5’-TATATAGGAAAGATATCCGGGTGAACTTCGGAATTTTGTCGCGATTCGAGGTTTTAG AGCTAGAAATAGCAAG-3’ and 5’-ATTTTAACTTGCTATTTCTAGCTCTAAAACAAGGATGCTTGTACTCGACTCGACGTTAAATTGAAAATAGGTC-3’. The backbone and insert were combined by Gibson Assembly Master Mix (New England Biolabs, E2611L). The gRNA plasmid was confirmed by sequencing and injected into transgenic embryos expressing Vasa-Cas9 and allowed to develop to adulthood (BestGene Inc., Chino Hills, USA). Each adult fly was individually crossed with a balancer stock to generate stocks of putative mutants. Genomic DNA was extracted from homozygous putative mutants of each stock for PCR amplification of *eIF4E-5* and sequenced using primers: 5’-GGTGATGACACTACTGACGC-3’ and 5’-AACGCCCAACAAACTGAAAC-3’ (The Centre for Applied Genomics, The Hospital for Sick Children, Toronto, Canada). This experiment was repeated twice; the initial round identified two different mutant alleles from the same founder parent (*eIF4E-5^B8a^* and *eIF4E-5^1B8b^*) and the second round identified a frame-shift allele (*eIF4E-5^D19a^*).

### Generation of genomic 3xFLAG-eIF4E-5 rescue construct and transgenic flies

The rescue construct consisted of the genomic DNA starting 359 bp upstream of the ATG, 69 bp 3xFLAG tag (5’-GACTACAAAGACCATGACGGTGATTATAAAGATCATGACATCGATTACAAGGATGACGATGACAAG-3’), 12 bp linker (5’-GGCAGCGAATTC-3’), all 791 bp of eIF4E-5 protein coding sequence including introns, and 362 bp downstream of the stop codon including the putative poly(A) site. The first three regions (5’ region, 3xFLAG, linker) was synthesized (BioBasic Inc., Markham, Canada) and subcloned into the pattB plasmid (Drosophila Genomics Research Center (DGRC), 1420, Bloomington, USA) using *Bam*HI (New England Biolabs, R3136S) and *Eco*RI (New England Biolabs, R3101S). The last two regions were PCR-amplified from genomic DNA with *Eco*RI and *NotI* (New England Biolabs, R3189S) added to the primers (5’-ATGACAAGGGCAGCGAATTCATGGCCAGTGCACAAGTG-3’and 5’-GTACCCTCGAGCCGCGGCCGCGCTTGAGTAGGCAATTACGAC-3’), and subcloned into pattB downstream and in-frame with the 5’ genomic region, 3xFLAG and linker. The pattB-3xFLAG-eIF4E-5 plasmid was confirmed by sequencing and integrated into the attP40 site on the second chromosome via PhiC31 integrase-mediated transgenesis by injection into *y^l^ M{3xP3-RFP-3xP3-GFP-vas-int.DM}ZH-2A w*; P{CaryP}attP40* embryos (BestGene Inc.).

### Generation of anti-eIF4E-5 antibodies

The full-length coding region of eIF4E-5 was PCR-amplified using the plasmid 4E5-pCR2.1 as a template (Hernández et al., 2005) and subcloned into pRSET (Invitrogen, V35120) to create an expression construct. The plasmid was transformed into *E. coli* BL21 (DE3) (Novagen, 71012) to produce a His6X fusion recombinant protein according to the manufacturer’s instructions. The fusion protein was purified using Ni-NTA beads under denaturing conditions (Thermo Fisher Scientific, R90101, Waltham, USA). Polyclonal anti-eIF4E-5 antibodies were raised in rabbit against this recombinant protein (Comparative Medicine and Animal Resources Centre, McGill University, Montreal, Canada). The sera were separated and NaN3 was added at a concentration of 0.02%.

### Squashed preparations of *Drosophila* testes

Testes were dissected from 0 to 2-day old males (unless otherwise stated) in cold testis isolation buffer (TIB) (Casal et al., 1990). Whole testes were mounted on polylysine coated slides in TIB and squashed with a coverslip (Polysine, P4681-001, Thermo Fisher Scientific). Live images for Fig. 3 were acquired on an upright Zeiss Axioplan 2E epifluorescence microscope equipped with a 20x phase-contrast objective and an Axiocam black and white CCD camera using Axiovision software (Carl Zeiss, Oberkochen, Germany). Live images for Fig. S3 were acquired on an inverted Leica DMi8 epifluorescence microscope equipped with a 20x phase-contrast objective and a Leica K5 camera using Thunder Imaging System. All images were uniformly processed for brightness and contrast using Adobe Photoshop (San Jose, USA).

### Immunohistochemistry on squashed testis preparations

Testes were prepared as for live microscopy (above) and were processed for immunofluorescence as previously described (Hime et al., 1996). In brief, after mounting on polylysine coated slides (Polysine, Thermo Fisher Scientific, P4681-001, Waltham, USA), samples were squashed with a coverslip and frozen in liquid nitrogen. Coverslips were removed with a razor blade, and samples were immediately chilled in 95% ethanol for at least 10 minutes. Samples were fixed in PBS with 4% paraformaldehyde (Electron Microscope Sciences, 15710, Hadfield, USA) for 7-10 minutes, permeabilized in PBS with 0.37% Triton X-100 and 0.3% sodium deoxycholate for 30 minutes, and blocked in PBS with 0.1% Triton X-100 and 5% bovine serum albumin (PBTB; Sigma-Aldrich, A3912-100G, St. Louis, USA). Samples were incubated at 4°C overnight with primary antibodies, then washed with PBTB three times for 5 minutes and once for 15 minutes and incubated for 1 hour at room temperature with secondary antibodies diluted in PBTB. Samples were then washed with PBTB once for 15 minutes, stained (when applicable) with rhodamine-phalloidin in PBTB (1:200; Invitrogen, R415) for 30 minutes, washed with PBS with 0.1% Triton X-100 (PBT) for 15 minutes, stained with 4’,6-diamidino-2-phenylindole (DAPI) in PBT (1:1000; VWR, 89139-054, Radnor, USA) for 10 minutes and washed with PBT twice for 15 minutes. Samples were mounted in ProLong Diamond Antifade Mountant (Molecular Probes, P36961, Eugene, USA), sealed with nail polish, and examined within 1-2 days. Fluorescence micrographs were acquired on a Nikon A1R inverted laser scanning confocal equipped with 10x, 20x, 40x, and 60x objectives, photomultiplier tube (PMT) detectors for DAPI channel, and gallium arsenide phosphide (GaAsP) PMT detectors for green and red channels using NIS Elements software (SickKids Imaging Facility, The Hospital for Sick Children, Toronto, Canada). All images were uniformly processed for brightness and contrast using Adobe Photoshop (San Jose, USA). Apart from Fig. 3H, all images from the same experiment were adjusted for brightness and contrast in an identical manner.

Primary antibodies used for immunofluorescence were rabbit anti-eIF4E-5 (1:500; #4524, this work), rabbit anti-caspase-3 (1:400; Asp175, Cell Signaling Technology, Danvers, USA), guinea pig anti-Soti (1:100; Kaplan et al., 2010; a kind gift of Eli Arama, Weitzmann Institute, Rohovot, Israel), mouse anti-Adducin 1B1 (1:20; Zaccai and Lipshitz, 1996; a kind gift of Howard Lipshitz, University of Toronto, Toronto, Canada) and mouse anti-FLAG (1:200; M2, Sigma-Aldrich, F1804). Secondary antibodies used for immunofluorescence were Alexa Fluor 488-conjugated anti-rabbit (1:1000; Invitrogen, A-11008), Alexa Fluor 488-conjugated anti-guinea pig (1:1000; Invitrogen, A-11073), or Alexa Fluor 568-conjugated anti-mouse IgG (1:1000; Invitrogen, A-31043).

### Quantifying spermatid cyst polarity defects

Testes were dissected and stained following the immunohistochemistry protocol described above. The percentage of testes with spermatid nuclei near the tip was recorded (Xu et al., 2014). For Fig. 7E-F, statistical analysis and graphing were performed using GraphPad Prism versions 8 for Macintosh, respectively (GraphPad Software). Differences observed between genotypes were analyzed with unpaired two-tailed Student *t*-test. Results were considered statistically significant at *p*<0.05.

### Immunofluorescence with single molecule RNA FISH

All solutions used were RNase free. Testes were dissected in 1xPBS (Invitrogen, AM9624) and fixed in 4% formaldehyde (Polysciences, Inc., 18814-10) in 1xPBS for 30 minutes, washed briefly in PBS, and permeabilized in 70% ethanol overnight at 4°C. Samples were then washed with 1xPBS and blocked for 30 minutes at 37°C in blocking buffer (1xPBS, 0.05% BSA [Invitrogen, Am2616], 50μg/mL yeast tRNA [Sigma-Aldrich, R8759], 10mM Vanadyl Ribonucleoside complex [New England Biolabs, S1402S], 0.2% Tween-20 [Sigma-Aldrich, P7949]). Samples were incubated with primary antibodies diluted in blocking buffer overnight at 4°C, washed with 1xPBS containing 0.2% Tween-20, re-blocked for 5 minutes at 37°C in blocking buffer, and incubated 4°C overnight in blocking buffer containing secondary antibodies. Testes were then washed with 1xPBS containing 0.2% Tween-20, and re-fixed for 10 minutes before being briefly rinsed with wash buffer (2x saline-sodium citrate [SSC, Invitrogen, AM9770], 10% formamide [Fisher Scientific, BP227]), and then hybridized overnight at 37°C in hybridization buffer (2xSSC, 10% dextran sulfate [Sigma-Aldrich, D8906], 1mg/mL yeast tRNA, 2mM Vanadyl Ribonucleoside complex, 0.5% BSA, 10% formamide). Following hybridization, samples were washed three times in wash buffer for 20 minutes each at 37°C and mounted in Vectashield with DAPI (Vector Laboratory, H-1200, Burlingame, USA). Images were acquired using an upright Leica Stellaris 8 confocal microscope with a 63X oil immersion objective lens (NA = 1.4) and processed using ImageJ software (National Institutes of Health, Bethesda, USA).

Primary antibodies were rabbit anti-eIF4E-5 (1:500; #4524, this work) or rabbit anti-FLAG (1:500; Invitrogen, PA1-984B), and secondary antibodies were Alexa Fluor 488-conjugated anti-rabbit (1:200; Life Technologies, Carlsbad, USA). Fluorescently labeled probes were added to the hybridization buffer to a final concentration of 100nM. Probes against *kl-3, soti,* and *eIF4E-5* mRNA were designed using the Stellaris® RNA FISH Probe Designer (Biosearch Technologies, Inc., Novato, USA) available online at www.biosearchtech.com/stellarisdesigner. Probe sets are listed in Table S1.

For smFISH alone, hybridization immediately followed the overnight incubation in 70% ethanol and short wash with wash buffer.

### Immunoblotting

Three methods were used for immunoblotting. For Fig. 1, approximately 20 pairs of testes per genotype were dissected in TIB with protease inhibitor cocktail (1:100; Halt, Thermo Fisher Scientific, 87786), then lysed in radioimmunoprecipitation assay (RIPA) buffer with protease inhibitors (150 mM NaCl, 50 mM Tris-HCl pH 8.0, 1% Nonidet P-40, 0.5% sodium deoxycholate, 0.1% SDS). NuPAGE LDS Sample Buffer was added, and samples were boiled for 10 minutes at 98°C (Invitrogen, NP0007). Proteins were run on gradient pre-cast SDS polyacrylamide gels (8-16%, ExpressPlu, GenScript, M81610, Piscataway, USA) before being transferred to nitrocellulose membranes (0.45μm, Amersham Protran, GE Healthcare Life Sciences, 10600020 Chicago, USA) with NuPAGE Transfer Buffer (Invitrogen, NP0006). Membranes were rinsed in TBST (Tris-buffered saline with 0.05% Tween-20), blocked in TBST containing 5% nonfat milk, and incubated overnight at 4°C with primary antibodies diluted in TBST containing 5% nonfat milk. Membranes were washed with TBST and incubated with secondary antibodies diluted in TBST containing 1% nonfat milk. Membranes were washed with TBST, and detection was performed using Novex ECL Chemiluminescent Substrate Reagents Kit (Invitrogen, WP20005). Primary antibodies used were rabbit anti-eIF4E-5 (1:5000; #4524, this work) and mouse anti-α-tubulin (5ug/mL; AA4.3, Developmental Studies Hybridoma Bank, Iowa City, USA). Secondary antibodies were HRP-conjugated anti-rabbit (1:10,000; Jackson ImmunoResearch Laboratories, 111-035-003, West Grove, USA) or anti-mouse IgG (1:10,000; Jackson ImmunoResearch Laboratories, 715-035-150).

For Fig. 6, testes (40 pairs/sample) were dissected in Schneider’s medium (Gibco, 2172-0024) at room temperature within 30 minutes, the medium was removed, and samples were frozen at −80°C until use. Tissues were then lysed in equal volumes of 2xLaemmli Sample Buffer (Bio-Rad Laboratories, 1610737, Hercules, USA) + βME (Bio-Rad Laboratories, 1610710, Hercules, USA) and equal volumes were run on a NuPAGE Tris-Acetate gel (3-8%, 1.5mm, Invitrogen, EA0378BOX) with Tris-Acetate SDS Running Buffer (Invitrogen, LA0041) before being transferred onto polyvinylidene fluoride (PVDF) membrane (Bio-Rad Laboratories, 1620177, Hercules, USA) using NuPAGE transfer buffer (Invitrogen, NP0006) without added methanol. Membranes were blocked in 1xTBST (0.1% Tween-20) containing 5% nonfat milk (Bio-Rad, 1706404, Hercules, USA), followed by incubation with primary antibodies diluted in 1X TBST containing 5% nonfat milk. Membranes were then washed with 1xTBST, followed by incubation with secondary antibodies diluted in 1xTBST containing 5% nonfat milk. After washing with 1xTBST, detection was performed using the Pierce® ECL Western Blotting Substrate enhanced chemiluminescence system (Thermo Fisher Scientific, 32106). Primary antibodies used were mouse anti-α-tubulin (1:2,000; clone DM1a, Sigma-Aldrich) and mouse anti-FLAG (1:2,500; M2, Sigma-Aldrich). Secondary antibody was HRP-conjugated anti-mouse IgG (1:10,000; Abcam, ab6789, Cambridge, UK).

For Fig. S6, immunoblotting was carried out as described for Fig. 6 except that samples were run on a Novex Tris-Glycine gel (14%, 1mm, Invitrogen, XP00140BOX) with running buffer (25mM Tris base, 192mM glycine, 0.1% SDS) and transferred to the PVDF membrane using transfer buffer (25mM Tris base, 192mM glycine, 20% methanol). Primary antibodies used were mouse anti-α-tubulin (1:2,000; clone DM1a, Sigma-Aldrich) and rabbit anti-eIF4E-5 (1:5000; #4524, this work). Secondary antibodies were HRP-conjugated anti-mouse IgG (1:10,000; Abcam, ab6789, Cambridge, UK) and HRP-conjugated anti-rabbit IgG (1:10,000; Abcam, ab6721, Cambridge, UK).

### Yeast two-hybrid system

A cDNA encoding *Drosophila* eIF4E-5 (CG8277) was PCR-amplified using the plasmid 4E5-pCR2.1 as a template (Hernández et al., 2005) and subcloned into the vector pOAD (“prey” vector; Cagney et al., 2000) in-frame with the activator domain sequence of GAL4 to generate the construct eIF4E-5-AD. *Drosophila* GRB10-interacting GYF (glycine-tyrosine-phenylalanine domain) protein (GIGYF, CG11148; Russica et al., 2019; a kind gift of Catia Igreja, Max Planck for Developmental Biology, Tübingen, Germany), CUP (CG11181; Nelson et al., 2004; Zappavigna, et al., 2004; a kind gift of Nancy Standart, Cambridge University, Cambridge, UK), eIF4E transporter (4E-T, CG32016), eIF4G (CG10811; Hernández et al. 1998), eIF4G-2 (CG10192; Baker and Fuller 2007), and 4E-BP (CG8846; Miron et al., 2001) cDNAs were subcloned into the pOBD2 vector (“bait” vector; Cagney et al., 2000) in-frame with the DNA-binding domain sequence of GAL4 to create the respective plasmids pGIGYF-BD, pCUP-BD, p4E-T-BD, peIF4G-2 (313-1164)-BD, peIF4G-BD and p4E-BP-BD.

Interactions between proteins expressed as “prey” or “bait” fusions were detected following a yeast interaction-mating method using the strains PJ69-4a and PJ69-4α (Cagney et al., 2000). Diploid cells containing both bait and pray plasmids were grown in –Trp, –Leu selective media (Clontech, 630417, Mountain View, USA) and shown as controls for growth. Protein interactions were detected by replica-plating diploid cells onto –Trp, –Leu, –Ade (20 ug/mL L-His HCl monohydrate (A-9126, Sigma) added to –Trp, –Leu, –Ade, –His (630428, Clontech) or –Trp, –Leu, –His (630419, Clontech) selective media +3 mM, 10 mM or 30 mM 3-amino-1,2,4-triazole (3AT, Sigma-Aldrich). Growth was scored after 4 days of growth at 30°C.

## Acknowledgments

The authors are grateful to Brill lab members Jonathan Ma, Nigel Giffiths, Lacramioara Fabian, Alind Gupta and Yonit Bernstein for their support and assistance with experimental methods. We thank Eli Arama and Howard Lipshitz for antibodies. We also thank Howard Lipshitz, Craig Smibert and James Ellis for helpful discussions, Helen White-Cooper for insightful comments on the manuscript, Paul Paroutis and Kimberly Lau of the SickKids Imaging facility for advice on imaging, and the Bloomington Drosophila Stock Center for fly stocks.

## Author contributions

Conceptualization, L.S., J.A.B.; Methodology, L.S., J.M.F., B.L.F., H.H., G.M., Y.Q., V.L., E.H., L.C., G.P., G.H., P.L., J.A.B.; Validation, L.S., J.M.F., B.L.F., G.M., G.H., J.A.B.; Formal analysis, L.S., J.M.F., B.L.F., J.A.B.; Investigation, L.S., J.M.F., B.L.F., G.M., V.L., E.H., L.C., J.A.B.; Resources, H.H., P.L.; Writing – original draft, L.S., J.A.B.; Writing – review and editing, all authors; Visualization, L.S., J.M.F., B.L.F., G.M., G.H., J.A.B.; Supervision, L.S., G.P., G.H., P.L., J.A.B; Project administration, L.S., J.A.B.; Funding acquisition, G.H., P.L., J.A.B.

---

**Fig. S1.**
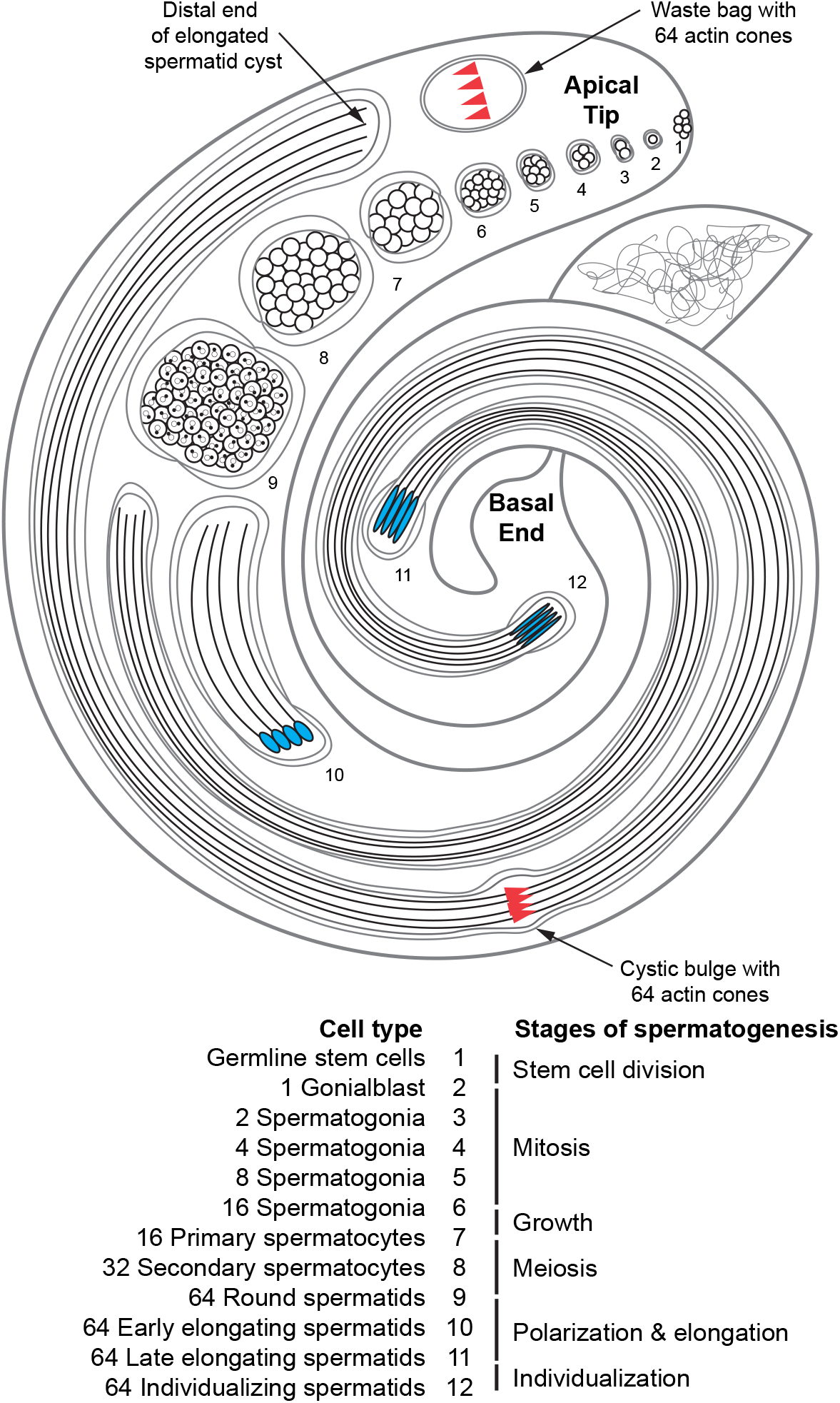
Stages of spermatogenesis are organized in a spatiotemporal manner within the *Drosophila* testis. Developing germline cells are enclosed by somatic cyst cells throughout spermatogenesis. Cysts undergoing elongation and individualization have 64 spermatids but only four are shown in this schematic for simplicity.

**Fig. S2.**
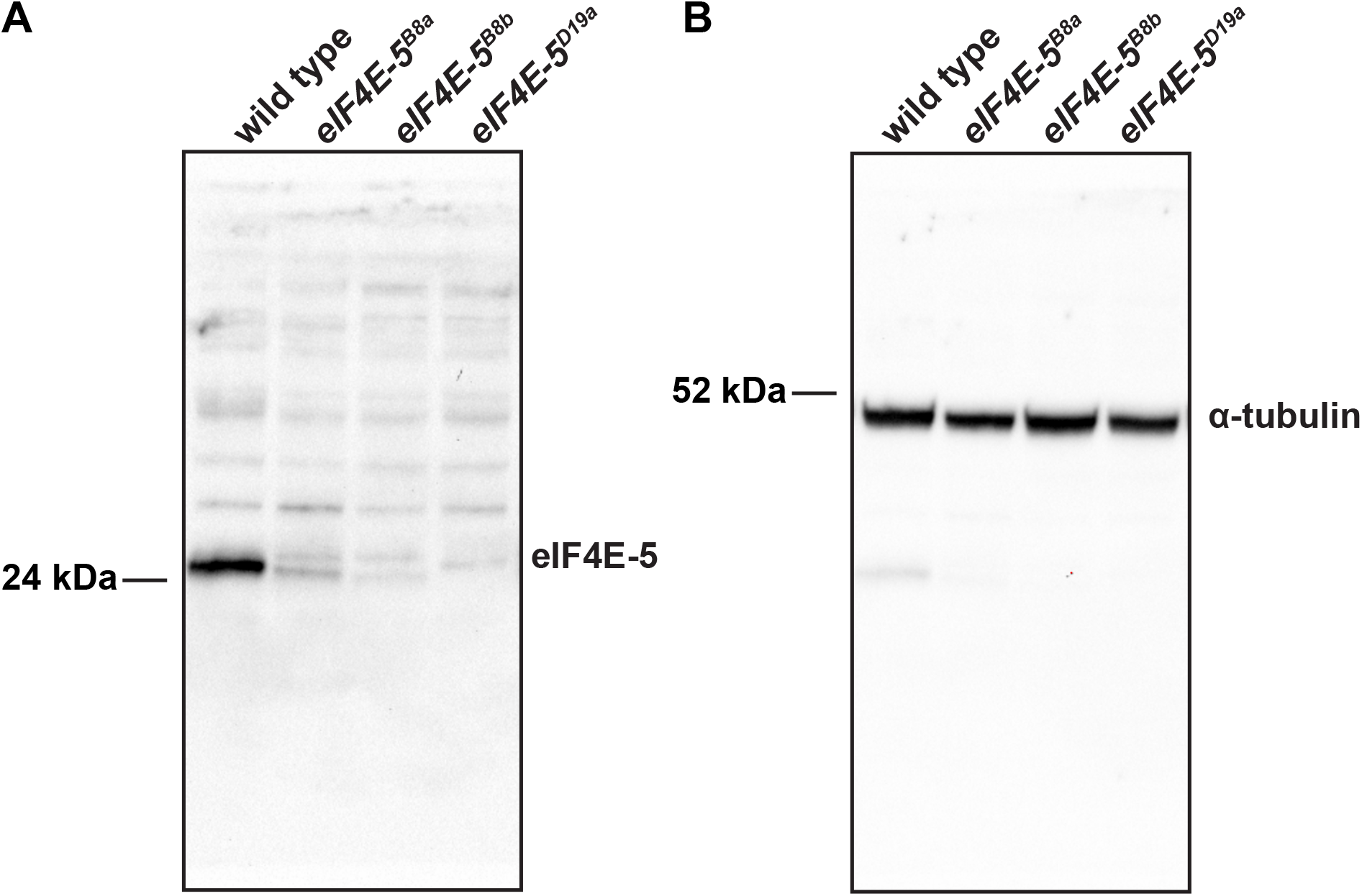
eIF4E-5 is reduced in *eIF4E-5* mutants. Whole immunoblots of total testis extracts probed with anti-eIF4E-5 (A) or anti-α-tubulin (B) reveal reduced (*eIF4E-5^B8a^* and *eIF4E-5^B8b^*) or undetectable (*eIF4E-5^D19^*) levels of eIF4E-5 protein while levels of tubulin loading control remained the same.

**Fig. S3.**
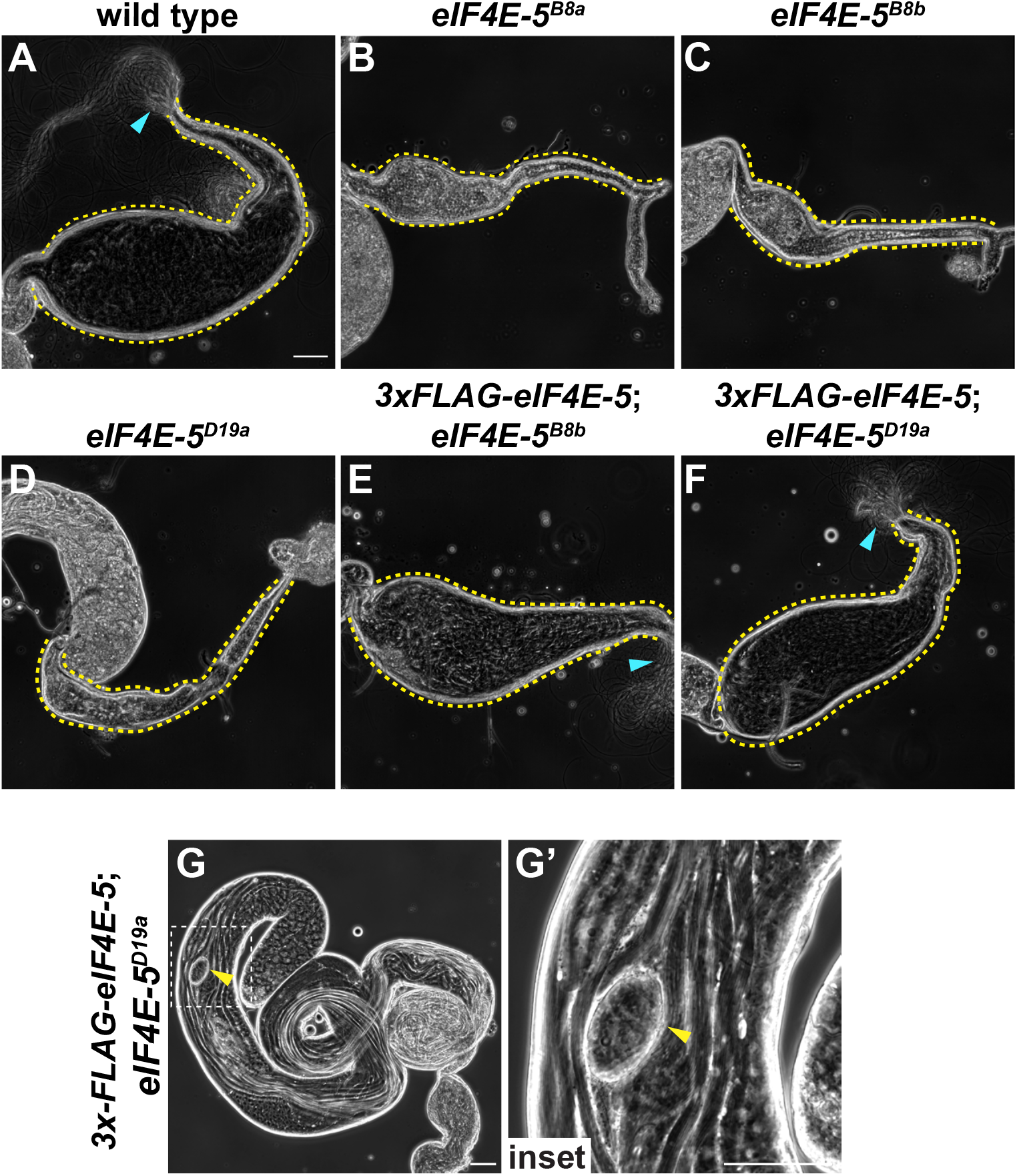
3xFLAG-eIF4E-5 transgene restores formation of mature sperm and waste bags in *eIF4E-5* mutants. Phase-contrast micrographs. **(A-F)** 7-day old wild type (A), *eIF4E-5^B8a^* (B, *eIF4E-5^B8b^* (C), *eIF4E-5^D19a^* (D), *3xFLAG-eIF4E-5;eIF4E-5^B8b^* (E), or *3xFLAG-eIF4E-5;eIF4E-5^D19a^* (F) revealing accumulation of mature sperm in seminal vesicles in mutants carrying 3xFLAG-eIF4E-5 transgene. Scale bar: 50 μm. **(G)** 3-day old *3xFLAG-eIF4E-5;eIF4E-5^D19a^* testis revealing presence of waste bag (yellow arrowheads). Scale bars: 50 μm.

**Fig. S4.**
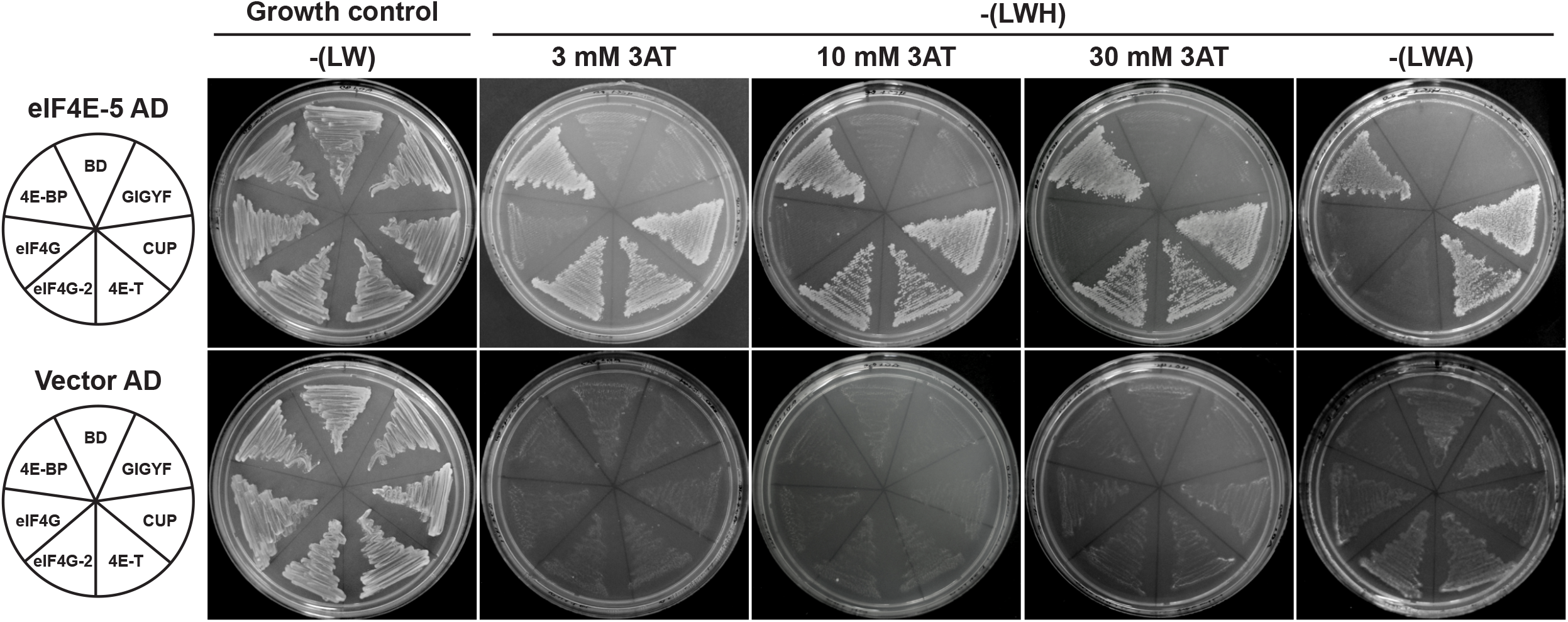
eIF4E-5 interacts with multiple translational regulators in yeast two-hybrid assays. eIF4E-5 interacts with Cup, 4E-T, eIF4G-2, and 4EBP in yeast two-hybrid assays, as revealed by growth on selective medium. Empty vectors (pOAD and pOBD2) were used as negative controls. L, leucine; W, tryptophan; A, adenine; H, histidine; 3AT, 3-amino-1,2,4-triazole.

**Fig. S5.**
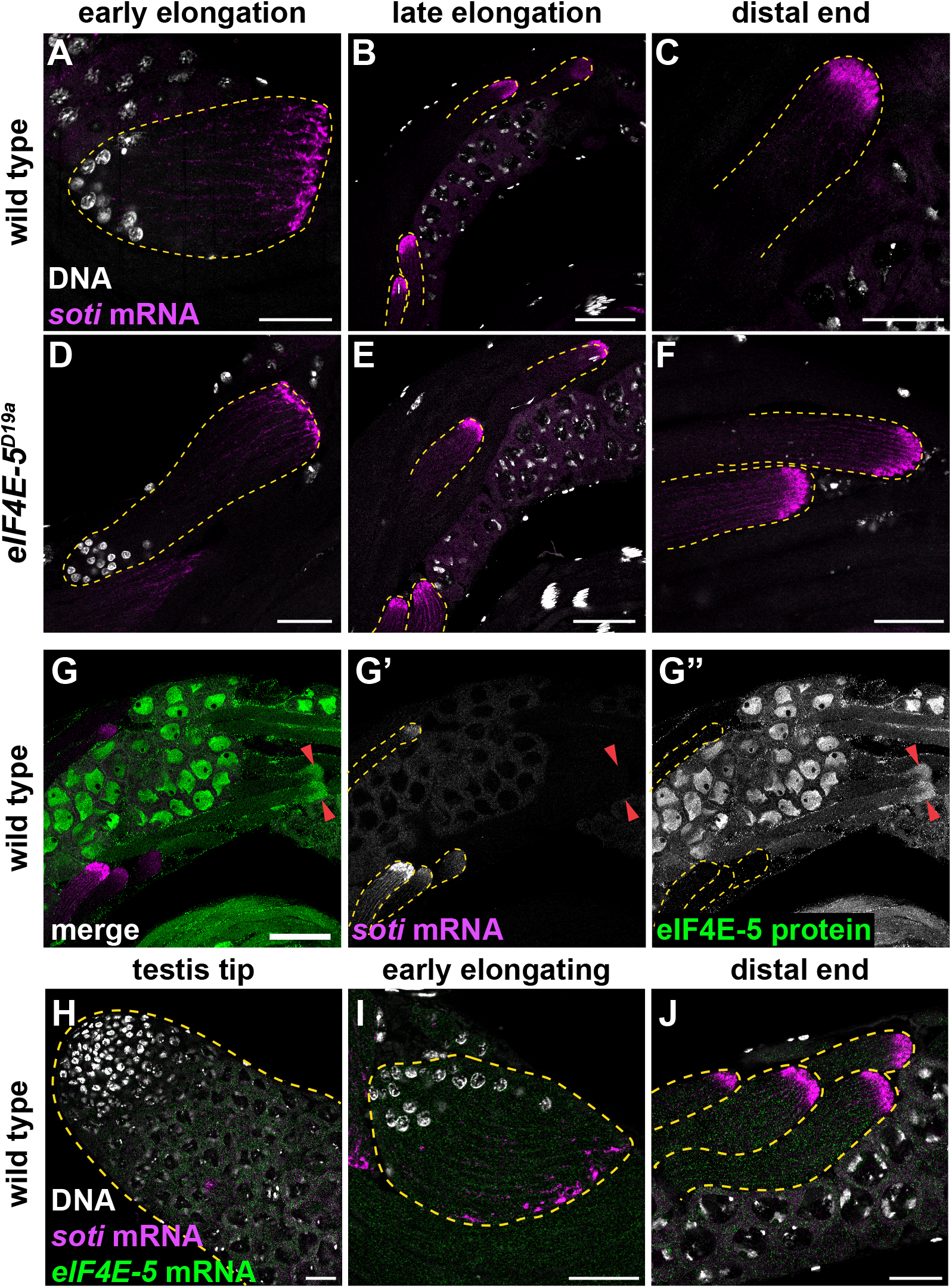
eIF4E-5 is dispensable for expression and localization of *soti* mRNA. **(A-F)** Laser-scanning confocal fluorescence micrographs of wild-type (A-C) and *eIF4E-5^D19a^* (D-F) whole adult testes probed for *soti* mRNA (magenta) and DNA (white). Expression and localization of *soti* mRNA at the distal ends of elongated cysts (cysts outlined by dotted yellow lines) appear similar in wild type and *eIF4E-5* mutants. Scale bars: 25 μm (A, D), 50 μm (B-C, E-F). **(G)** Laser-scanning confocal fluorescence micrographs of a wild-type whole adult testis probed for *soti* mRNA (magenta) and endogenous eIF4E-5 (green). *soti* mRNA (cysts outlined by dotted yellow lines) does not colocalize with eIF4E-5 at the distal ends of elongated spermatid cysts (red arrowheads). Scale bar: 50 μm. **(H-J)** Laser-scanning confocal fluorescence micrographs of wild-type whole adult testes probed for *soti* mRNA (magenta) *eIF4E-5* mRNA (green) and stained for DNA (white). Expression and localization of *eIF4E-5* mRNA is diffuse during early and late stages of spermatogenesis. Scale bar: 25 μm.

**Fig. S6.**
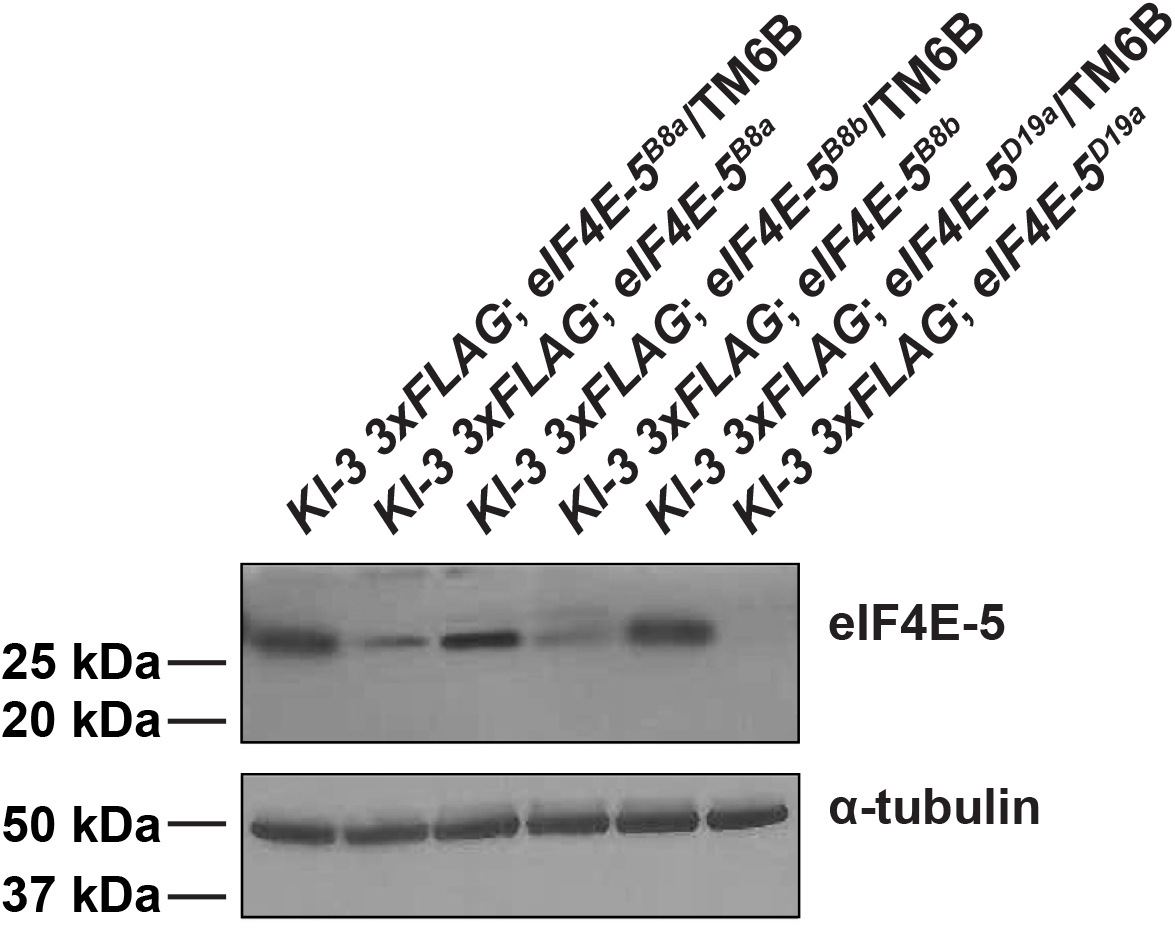
eIF4E-5 levels remain reduced in *Kl-3 3xFLAG;eIF4E-5* mutants. Immunoblots of whole testis extracts revealing eIF4E-5 levels in the indicated genotypes expressing endogenously tagged Kl-3 3xFLAG. eIF4E-5 protein levels are reduced in *eIF4E-5* mutants.

**Table S1.**
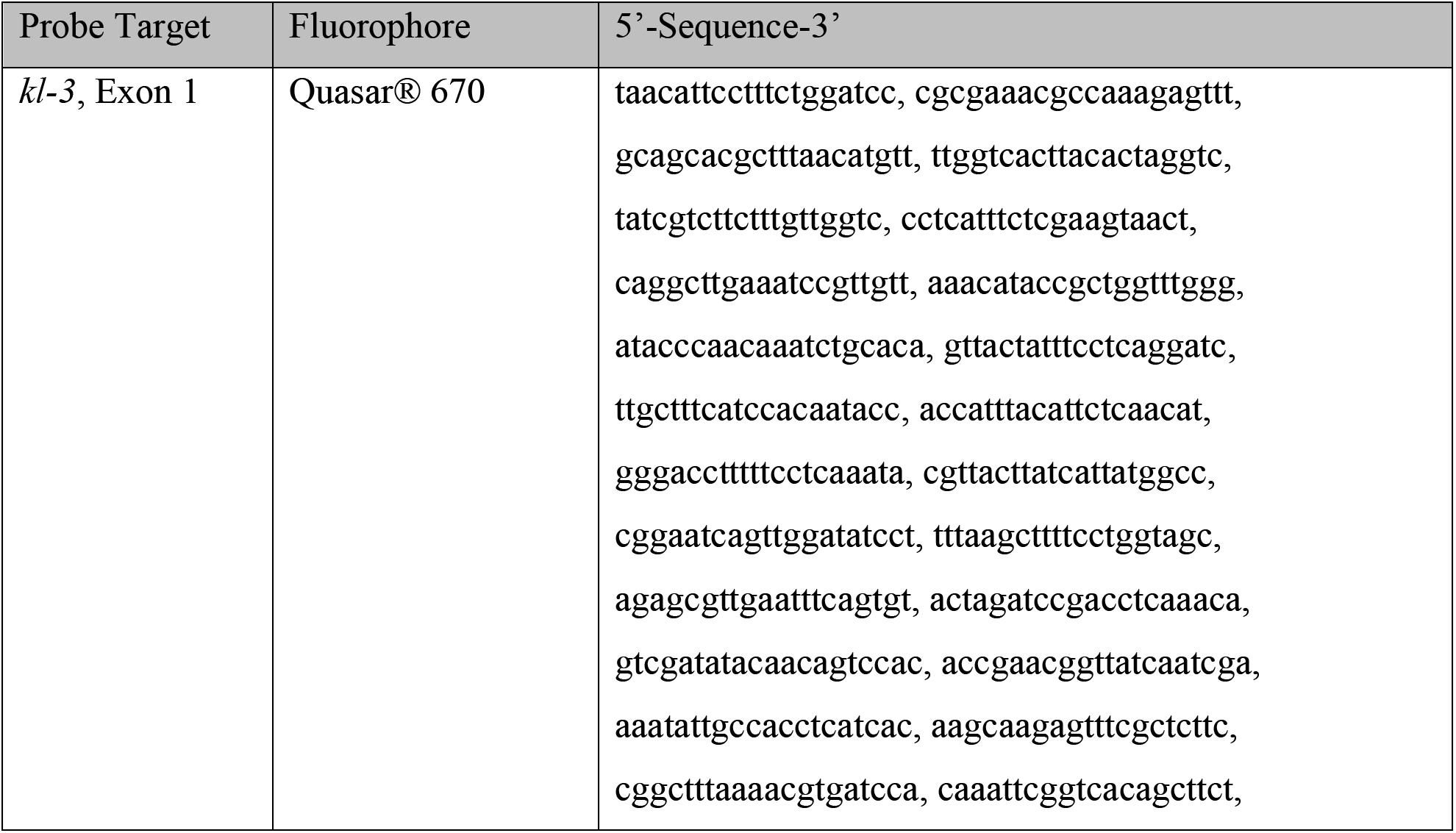
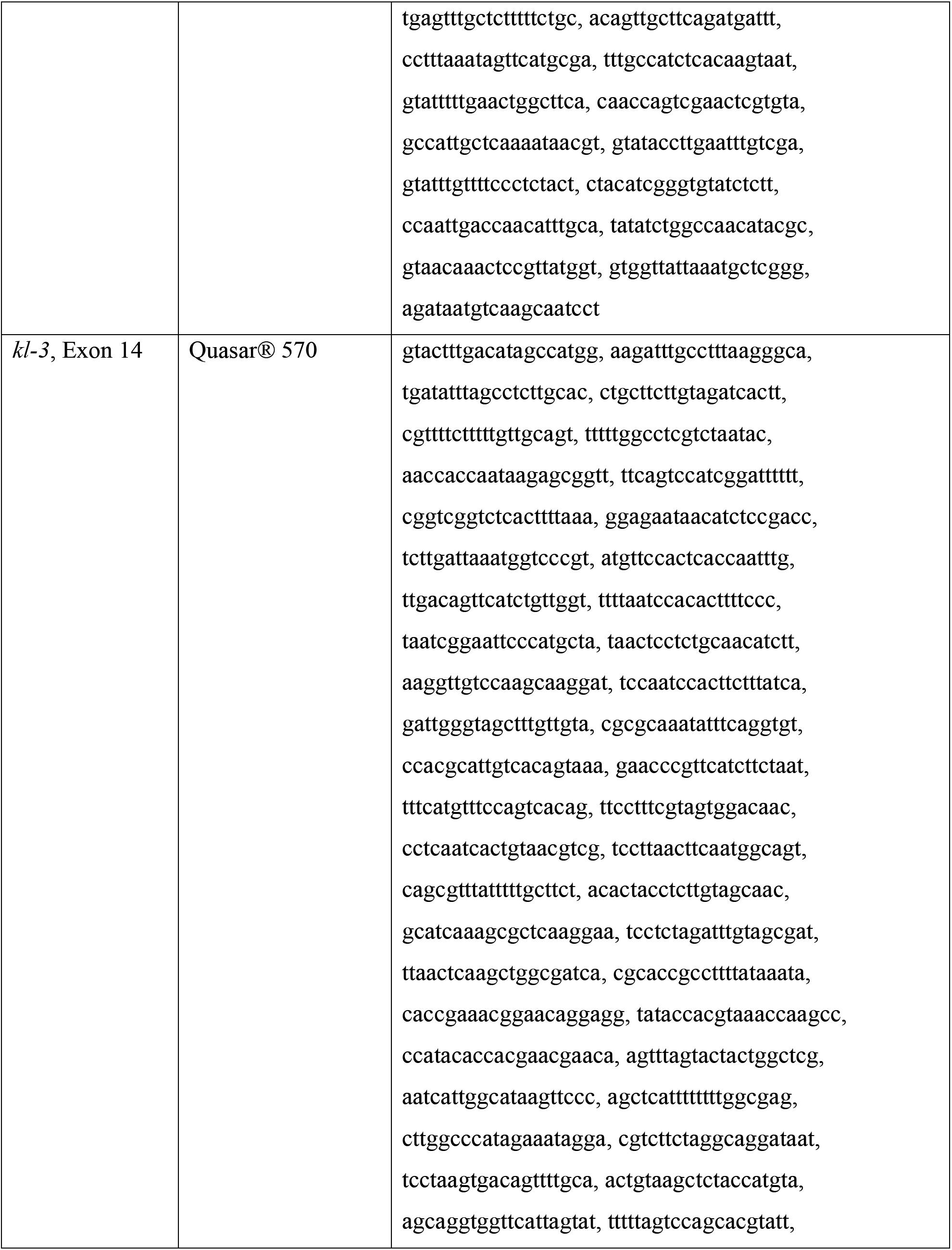
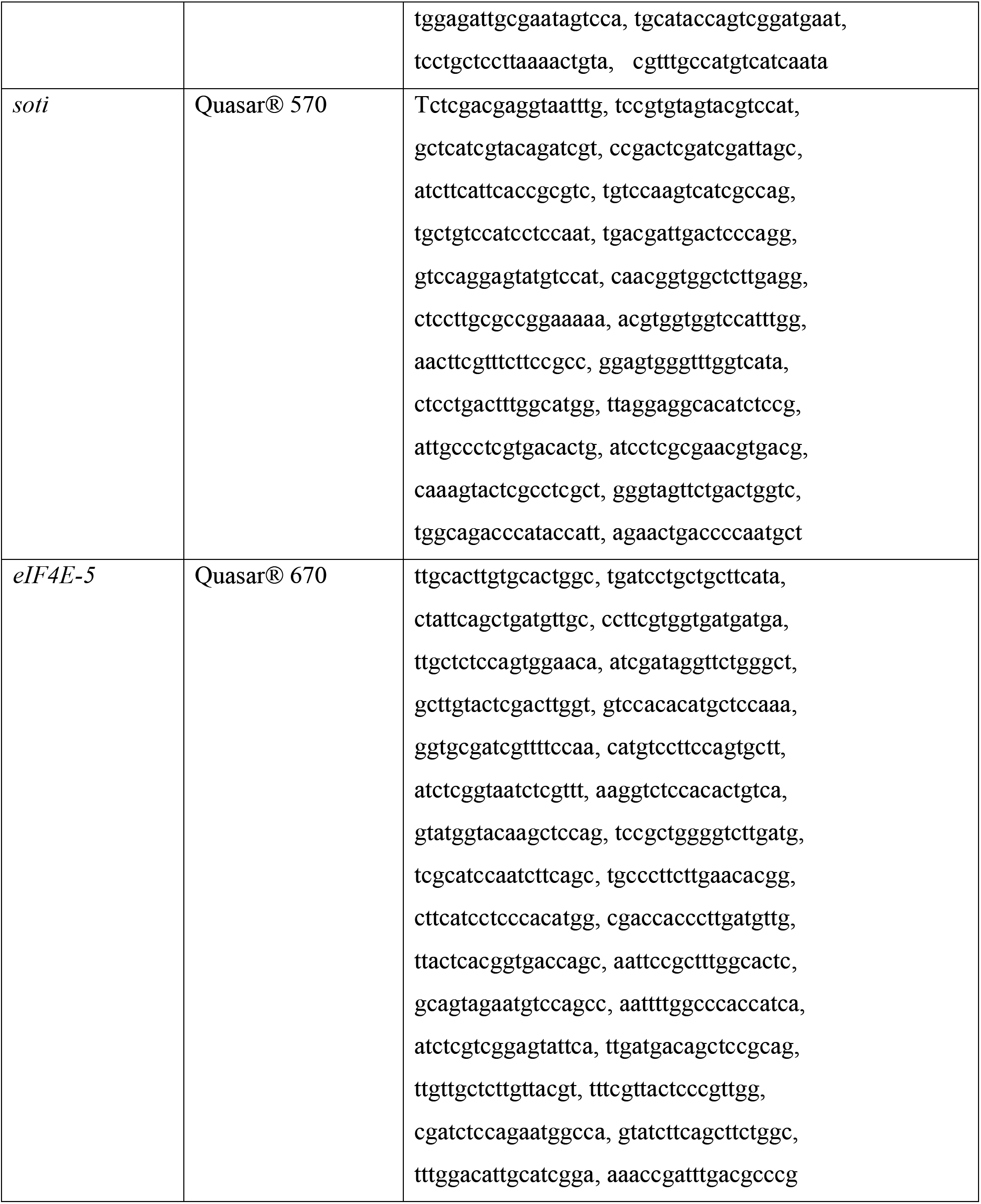
Probes for RNA FISH.

